# Viewing stomata in action: Autonomous *in planta* imaging of individual stomatal movement

**DOI:** 10.1101/2024.03.13.584774

**Authors:** Tomas E. van den Berg, Remco G. P. Sanders, Elias Kaiser, Jurriaan Schmitz

## Abstract

Stomata regulate plant gas exchange under changing environments, but observations of the dynamics of single stomata *in planta* are sparse. We developed a compact microscope system that can measure the kinetics of tens of stomata *in planta* simultaneously, with sub-minute time resolution. Dark field imaging with green light was used to create 3D stacks from which 2D surface projection were constructed to resolve stomatal apertures within the field of view.

Stomatal dynamics of *Chrysanthemum morifolium (*Chrysanthemum) and *Zea Mays* (Maize) under dynamically changing light intensity were categorized, and a kinetic model was fitted to the data for quantitative comparison. In addition, we also resolved dynamics of the surface position of the leaf, related to dynamics of leaf thickness or bending.

Maize stomata oscillated frequently between open and closed states under constant growth light and these oscillating stomata responded faster to changes in light than non-oscillating stomata at the same aperture.

The slow closure of Chrysanthemum stomata reduced water use efficiency (WUE). Over 50% showed delayed or partial closure, leading to unnecessarily large apertures after reduced light. Stomata with larger apertures had more lag and similar closure speeds compared to those with smaller apertures and lag, further reducing WUE. In contrast, maize stomata with larger apertures closed faster, with no lag.

In conclusion, our new system enables fine mapping of the heterogeneity of movement in neighboring stomata, providing new insights on the relations between stomatal dimensions, relative position and aperture changes under fluctuating light intensity.

## Introduction

Due to climate-change driven increases in air temperature and Vapor Pressure Deficit (VPD) (Novick *et al*., 2016) and decreased rainfall in many agricultural areas (Dore, 2005), future crop yields are coming under pressure. Adaptation of agriculture is therefore essential to maintain food security for an increasing world population (Anderson, Bayer and Edwards, 2020). A pillar for agricultural adaptation is the development of varieties with increased water use efficiency (WUE). To achieve this, we need to thoroughly understand the role that stomata play in plant WUE, which is often defined as the rate of photosynthesis divided by the rate of transpiration (Bertolino, Caine and Gray, 2019; Buckley, 2019; Lawson and Vialet-Chabrand, 2019; Nadal and Flexas, 2019), to breed for stomatal traits underlying high WUE.

Stomata are the microscopic pores on plant leaves that regulate their gas exchange, by dynamically opening and closing in response to environmental and intrinsic stimuli (Lawson and Matthews, 2020). Their dynamic behavior (Lawson and Vialet-Chabrand, 2019), morphology and density (Bertolino, Caine and Gray, 2019; Duursma *et al*., 2019) are important for WUE. Stomatal conductance to CO_2_ diffusion into the leaf facilitates photosynthetic CO_2_ fixation in the mesophyll, and stomatal movement broadly aims to balance the CO_2_ taken up by photosynthesis with the water vapor lost through transpiration. Light intensity changes of ~25-50 fold on single leaves are frequent in the crop growth environment (Kaiser, Morales and Harbinson, 2018). When the light intensity drops, the photosynthetic demand for CO_2_ decreases near-instantaneously, while stomatal closure proceeds much more slowly. Hence, stomata that at that moment are open more than strictly necessary for CO_2_ demand, waste water through unnecessary transpiration. Fast-closing stomata are thus more water use efficient (Papanatsiou *et al*., 2019; Horaruang *et al*., 2022).

An increase in light intensity quickly causes photosynthetic demand for CO_2_ to increase, as biochemical limitations are lifted in the first minutes of photosynthetic induction (Sakoda *et al*., 2021). To meet this demand for CO_2_, stomata need to open to increase the rate of diffusion of CO_2_, leading to a transient limitation of photosynthesis while opening. Transient limitations of photosynthesis in shade-sun transitions may cost an estimated 10-40% of potential crop CO_2_ assimilation (Long *et al*., 2022).

Stomatal dynamics are most frequently studied using leaf-level gas exchange measurements and are then related to static microscopic observations of e.g. leaf epidermal peels to stomatal density and morphology. Alternatively, dynamic behaviour of individual stomata has been studied on epidermal peels, missing the interaction with the mesophyll (Weyers and Travis, 1981; Blatt, 1987; Pantaleno *et al*., 2024). Measurements of bulk stomatal behavior, such as those of leaf gas exchange, hide the variation between individual stomata (Kaiser and Paoletti, 2014), and studies that have resolved individual stomatal dynamics *in planta* are relatively rare (Kaiser and Kappen, 1997, 2000, 2001; Kaiser, 2009; Grantz, Zinsmeister and Burkhardt, 2018). Moreover, minimizing the boundary layer of still air surrounding the leaf, as is common in gas exchange measurements (Busch *et al*., 2024), limits the study of stomatal behavior under frequently occurring natural conditions, when a significant boundary layer is present (Pattey *et al*., 1991; Goldstein *et al*., 1995). Information is lacking on how local morphology and density influence the dynamics of individual stomata. Such characteristics could lead to a better understanding of stomatal control and breeding targets for WUE and yield (Haworth *et al*., 2021). Additionally, it is essential to study such characteristics in the field, e.g. to understand how plants with altered stomatal characteristics respond to multiple stresses in different developmental phases (Bertolino, Caine and Gray, 2019).

Here, we describe a newly developed portable microscope, which can measure the opening and closure of tens of individual stomata simultaneously in the growth environment. Our method innovates on previous methods by selective use of green light for imaging and blue and red light as actinic light. In addition, we imaged the entire field of view (FOV) of the surface of the leaf, by creating large image stacks that were used to create leaf surface projections, in contrast to autofocus onto single pores (Kaiser and Kappen, 1997; Grantz, Zinsmeister and Burkhardt, 2018). This enabled us to relate stomatal dynamics to characteristics such as ‘the Voronoi area’, the leaf area with the shortest path length to a given stoma. Acquisition and analyses were largely automated, facilitating easy measurements. We demonstrated our method on leaves of Chrysanthemum, a greenhouse crop with elliptic stomata whose stomatal behavior can limit vase life (Fanourakis *et al*., 2021) and the rate of photosynthetic induction (Zhang *et al*., 2022) and the staple crop Maize with dumbbell stomata.

## Materials and Methods

### Microscope

The design of our microscope focused on obtaining images with sub-micrometer resolution at different focus planes to resolve the 3D surface of the leaf with sub-minute time resolution. Additionally, the design aimed to be minimally invasive to the leaf’s microclimate.

We achieved high optical resolution with high quality optics (plan APO series objectives, Mitutoyo, Japan and TTL200-S8 tube lens, Thorlabs, United States), verified with a calibration target (micro V2, Opto, Germany). The 20x objective (NA 0.42) enabled imaging of ~0.45 mm^2^ (722×625 µm) of the leaf’s surface. High time resolution of stacks of 100 images spaced 1 µm apart on the axis perpendicular to the imaging plane (z-axis; below the 1.6 µm depth of focus of the microscope objective) was achieved with a fast and accurate step motor (Z812, Thorlabs, United States, range of 12 mm, 0.2 µm precision), which was controlled with autofocus software and a CMOS camera (BFS-U3-244S8M-C; Teledyne-FLIR, Wilsonville, OR, USA). The high sensitivity of the camera and on-camera pixel binning (2×2) allowed us to achieve high quality images at a relatively low light intensity (50-75 µmol photons m^−2^ s^−1^) and short integration time (400 ± 100 ms). The use of long working distance objectives (20 mm) and leaf clips that were laser-cut from transparent polycarbonate with 2 mm neoprene cushions allowed us to image the leaf with low obstruction to air flow and light. Imaging made use of green light only (Effiring 525 nm; Effilux, Hürth-Efferen, Germany, for Chrysanthemum, SL-3500, PSI, Czech Republic, for Maize) by filtering light that passed through the objective with a bandpass filter (FB550-40, 550 ± 8 nm, FWHM 40 ± 8 nm; Thorlabs, Newton, NJ, USA). Monochromatic light was chosen, because it boosted image quality by limiting chromatic aberrations. Green light was selected because of its high reflectance by and transmission through the leaf, higher optical resolution compared to near-infrared radiation, and a maximum sensitivity of the camera in the green waveband. Optimization of imaging light was achieved before each automated measurement, by manually changing the emission angle and diffusivity (by changing the opacity of the window) of the imaging light. Blue-red actinic illumination was emitted by a custom-built LED lamp for Chrysanthemum (444/661 nm, 20/12 nm FWHM, 52/48%; Seven steps to heaven, the Netherlands) and by an RGB LED panel (SL-3500, PSI, Czech Republic) for Maize. Imaging could be done in brightfield or darkfield (with the green ring light above the leaf and blue-red light passing through the central hole of the green LED ring for Chrysanthemum). Maize and Chrysanthemum were imaged in darkfield. Both actinic and imaging light were controlled via LabVIEW (2018, National Instruments, Austin, TX, USA) and intensity calibrated at the leaf position with a photosynthetically active radiation (PAR) quantum sensor (Li-190R; Li-Cor Biosciences, Lincoln, NE, USA). Leaves were clamped in the microscope leaf holder that was equipped with embedded magnets at four contact points. The portable microscope was mounted on a tripod (Fig. 1). PAR intensity fluctuations during the measurement were recorded with a leaf clip holder (2020-B) connected to a portable fluorometer (MINI-PAM; Walz, Effeltrich, Germany).

**Figure 1.**
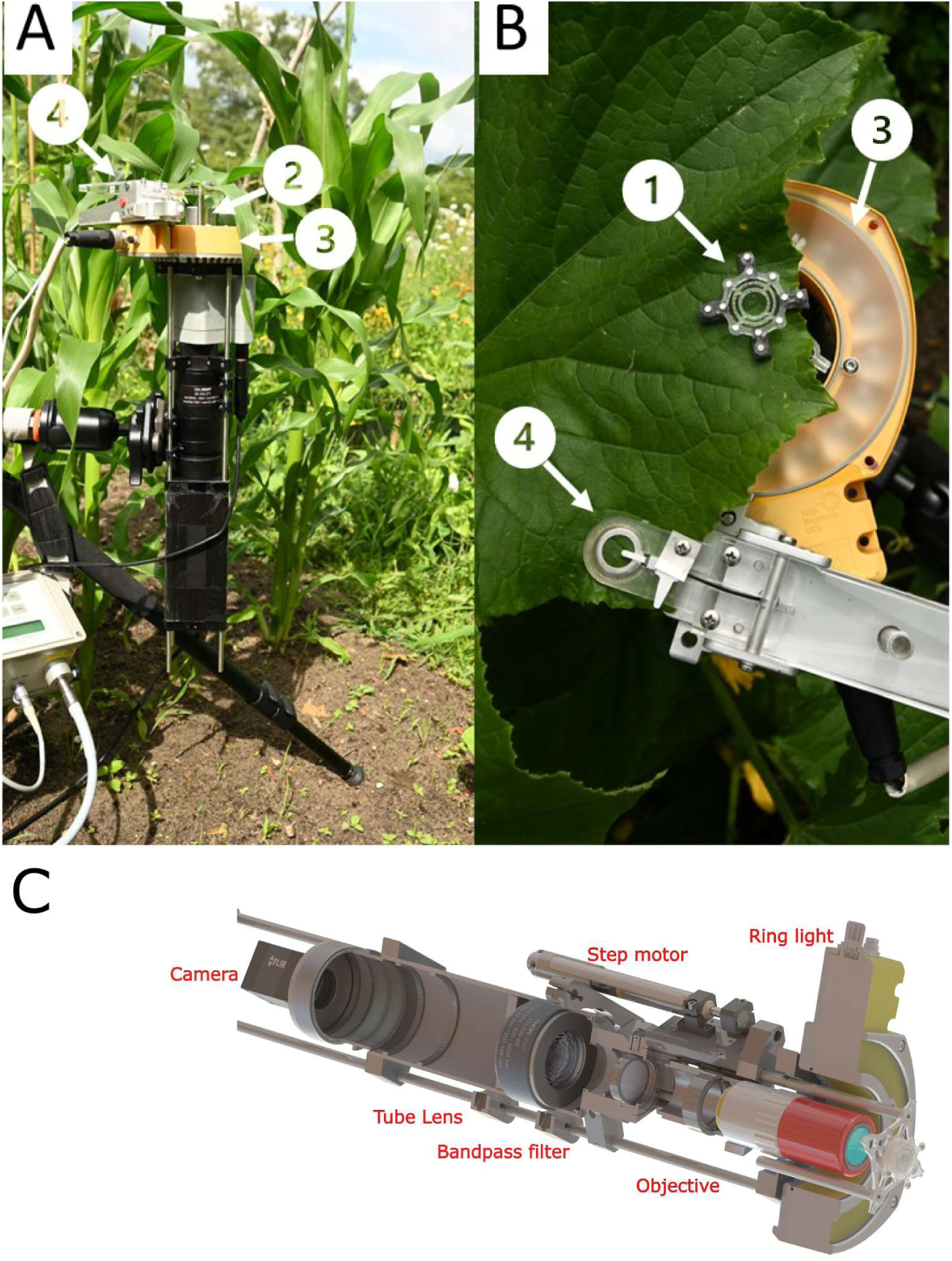
Overview of the leaf microscope. Example of the microscope A. Side view of the entire microscope with a leaf of maize (Maize) clamped in the holder. B. Top view of a leaf of cucumber (Cucumis sativa) clamped in the holder. The leaf clip (1) holds the leaf in place via magnets embedded in the polycarbonate framework. Neoprene cushions ensure a minimal effect of the clamping on the surface of the leaf. The microscope objective (2) can focus automatically, and the ring light (3) provides illumination for microscopy. The leaf clip from the mini-PAM (4) records PAR, ambient air temperature (A) or leaf temperature (B). C. Visualization of the layout with the most important components. Note that in this example, the green ring light is installed for brightfield imaging.

### Software

#### Acquisition

The data acquisition workflow was developed in Labview. Camera settings, number of images per stack with different focus planes on the z-axis (z-stack) and stack depth, imaging and light intensity and duration, as well as autofocus settings that determined the position of the stack’s center (methods: Roberts, Sobel, Gradient (Lthi *et al*., 2010)) were controlled via the interface in Labview. Full protocol files with imaging, light timing as well as light intensity settings can be loaded into the software with a graphical user interface for inspection.

#### Analyses

Acquired z-stack images, in 16bit TIFF format belonging to each z-stack, were processed in several stages (Fig. 2) using Fiji open-source software (Schindelin et al., 2012). Images were denoised (despeckle and outlier removal), normalized for intensity (enhance contrast, normalize) and aligned (SIFT linear stack alignment (Lowe, 2004)) (SM-Methods-Analyses). SM movie 1 is an example of a processed image stack. Next, the plugin “extended depth of field” (Forster *et al*., 2004) was used to generate a focus projection of the leaf surface, reducing the stack to a single image per timepoint (SM-Methods-Analyses). All timepoint focus projections were then stacked again to generate a video of the entire experiment that was again aligned to adjust for x-y axis movements of the leaf during the experiment (SIFT linear stack alignment). This stack was then cropped, to include only the area that was within view during the entire experiment, shadow corrected using the pseudo flat field correction in the BioVoxxel plugin (Brocher, 2015) and contrast enhanced using the contrast limited adaptive histogram equalization (CLAHE) algorithm (SM-Methods-Analyses) (Reza, 2004) (Fig. 3 A-C). SM movie 2 is an example of a processed video from an entire experiment, while SM movie 3 is a zoomed-in version of a single stoma. Auto thresholding was done in Percentile mode (Doyle, 1962) for Chrysanthemum (Fig. 3D) and in Berndsen mode (Sezgin and Sankur, 2004) for Maize, as it generated the best segmentation of open pores (SM-Methods-Analyses). SM movie 4 is SM movie 3 after thresholding. Stomata in the stack were manually selected with the elliptical (Chrysanthemum) or rectangular (Maize) selection tool and added in the region of interest (ROI) manager (ImageJ). Pore areas were then quantified via the ‘analyse particles’ menu for each stoma in the ROI manager, to quantify pore area and to generate masks for visual inspection of pore shapes for the entire experiment (Fig 3D). SM video 5 shows the masks generated from the stoma in SM videos 3 and 4.

**Figure 2.**
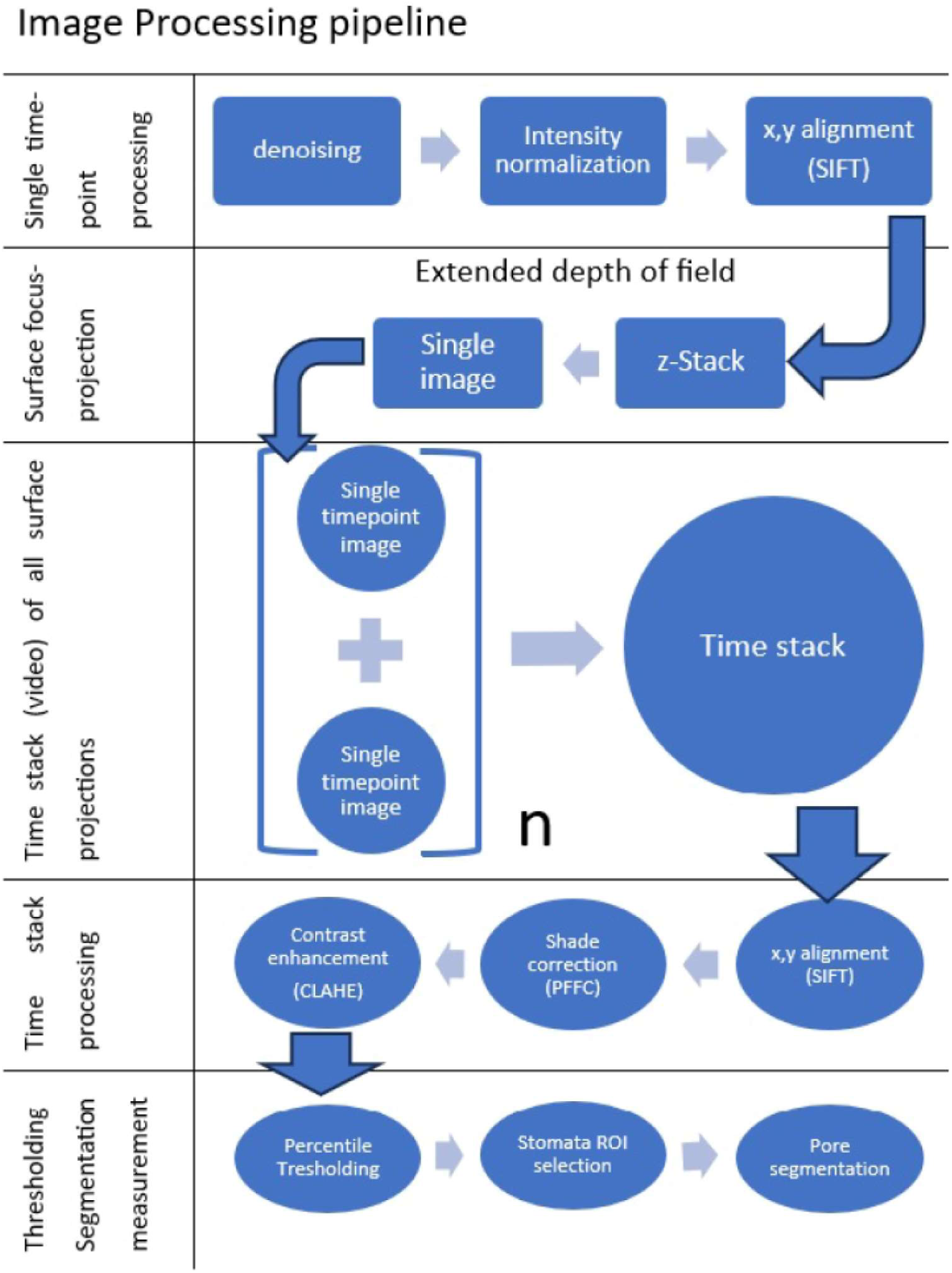
Flow chart of image processing pipeline. Single images belonging to an image stack were denoised and their contrast was enhanced by intensity normalization (SM-Methods-Analyses). Images with different z-axis position in a stack were then aligned to correct for minor movements of the leaf surface in x or y directions (surface parallel to the microscope’s objective). The extended depth of field plugin processed the stack to create a single focus projection of leaf surface (Fig. 3A). Each surface projection timepoint was added together in a video of the entire experiment. The images in these videos were again aligned to correct for movements in the x and y directions, shade corrected (Fig. 3B) and their contrast was enhanced (Fig. 3C). Automated thresholding of time stacks was done with Percentile (Chrysanthemum, Fig. 3D) or Berndsen mode (Maize). Stomatal positions were manually selected with the selection tool, the pore area was segmented, and quantified with the analyze particles menu.

**Figure 3.**
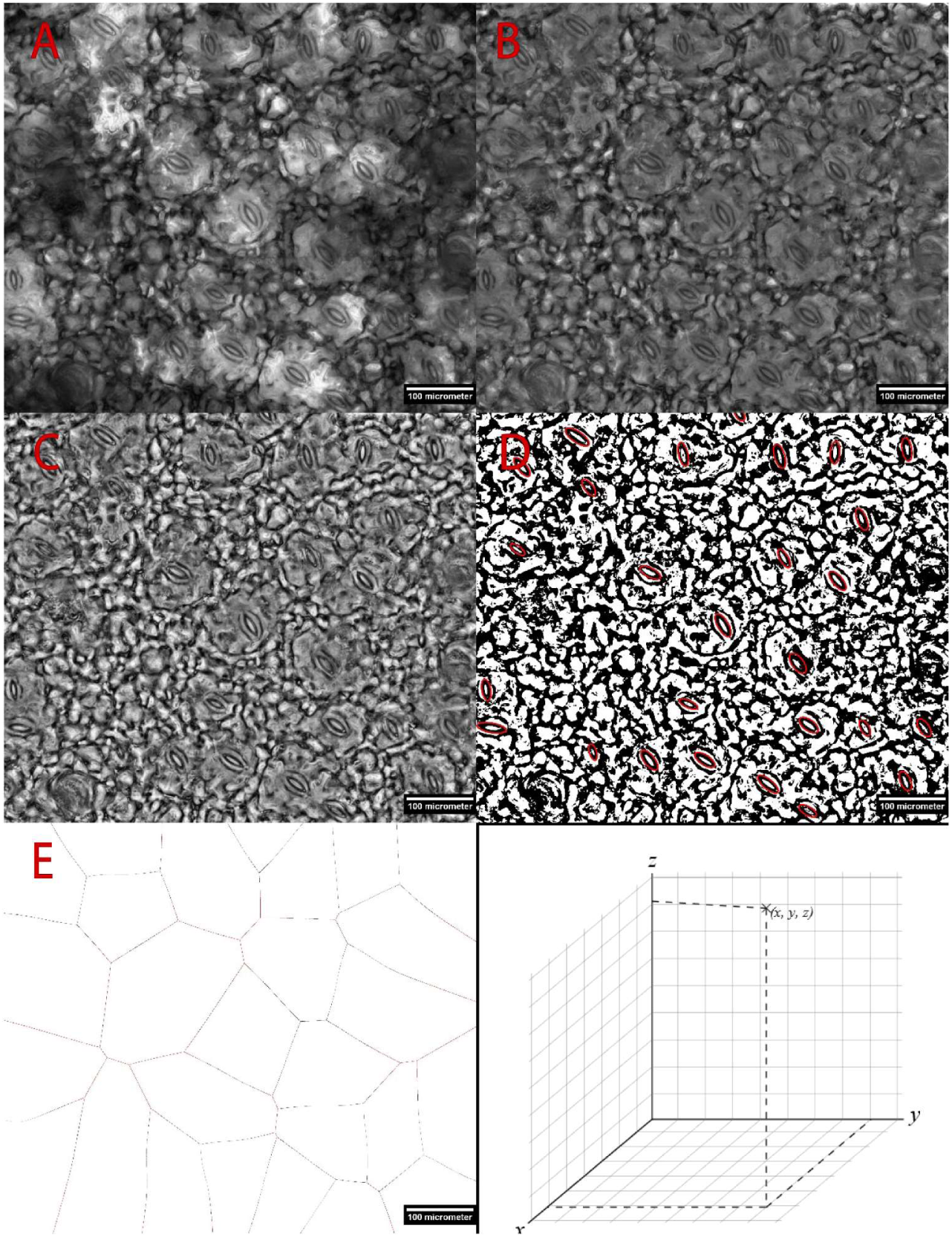
Example of time-stack processing with a single surface projection of Chrysanthemum. A. Surface projection output from extended depth of field (Forster et al., 2004). B. Image in A processed with pseudo flat field correction (PFFC, (Brocher, 2015). C. Image in B processed with contrast limited adaptive histogram equalization (CLAHE, (Reza, 2004). D. Image in C after thresholding with the Percentile method (Doyle, 1962). Red ROIs are drawn manually with the elliptical selection tool. E Voronoi diagram drawn based on the elliptical selections of stomata in D, only the Voronoi that are fully resolved are quantified. The scale bar indicates 100 micrometers.

Kinetics of stomatal aperture changes were loaded into Origin (OriginLab Corporation, Nothampton, MA, USA) and data of opening and closing were fitted separately with the model developed for kinetics of stomatal conductance (Vialet-Chabrand, Dreyer and Brendel, 2013):

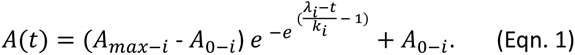

With A(t) the stomatal aperture at time t, A_max_ the maximum aperture at the final steady-state (SS), A_0-i_ the aperture at initial SS, λ the time lag of the response (min) and k the time constant (min), a measure of the rapidity of the response. For stomatal closure, A_min-d_ was defined as the final aperture at SS with A_0-d_, the aperture at the start of the closing response. k and λ were separately quantified for open and closing responses, using subscripts I (increase) and d (decrease), respectively

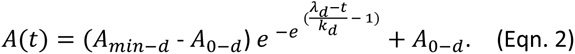

SM Fig. 1A provides an example of the fit of the kinetics for a single Chrysanthemum stoma.

Parameters that relate to SS pore aperture, such as k or A_max_ were only compared if the SS could be fitted with confidence (final aperture value in high light period (1000-1500 µmol photons m^−2^s^−1^) ≥ 90% offitted value of A_max_). In the 180-minute shade-sun-shade experiments in Chrysanthemum (SM Methods Lighting and Imaging Protocols), the steady state was not fitted with confidence, therefore k_i_, A_max-i_ and k_d_ were ignored in the analyses.

Because pore apertures in maize stomata frequently oscillated, we only fitted the first opening kinetics until a quasi-steady state (QSS) had been reached, to fit the starting aperture (A_0-i_), the opening delay (λ_i_) and the initial maximum opening speed (max dAdt_i_) (SM Fig 1B). The maximum aperture was not derived from fitting, but instead the maximum value obtained under high light intensity was used (SM Fig 1B). For stomata that opened to a complete steady state, we report all fit values. During closure, we again fitted only the first closing kinetics, until a QSS was reached for stomata that re-opened afterwards (SM Fig 1B). The minimum aperture after closing (A_min-d_) was taken from the fitted curves, because none of the stomata closed further.

Maximum opening and closing speeds (max dAdt_, I_ and _d_) were calculated by finding the maximum of the first derivative for the fitted curves within the time range of the fitted data (no extrapolation).

Some stomata were excluded from data analysis. Reasons to exclude stomata were: a) bad segmentation due to low local image quality, such as when a stoma shaded by a trichome, b) incomplete stomatal pore on the edge of an image, c) ‘false stomata’ stomata that never opened while image quality was not compromised, d) low-quality fit. Quality of fit was assessed by adjusted R-squared statistic of the fit (ideally close to 1), the structure in the residuals (ideally random noise without structure) and error estimates of the fitted parameters (ideally <10%, but no larger than 100%).

For Chrysanthemum 2% (3/124) of closing stomata were omitted from the kinetic analyses because the fit was bad (adjusted R-squared<0.8).

For maize, 17% (5/29) of opening and 21% (6/29) of closing kinetics of stomata were omitted from the kinetic analyses, as their fast kinetics could not be fitted accurately (40%-900% fit error) due to the measuring interval of the microscope. We report more on these stomata in the results.

### Categorization of kinetics

#### Chrysanthemum

Before a step increase in light intensity, stomata were categorized by fitting a straight line through the initial time course of pore aperture in low light. If the slope was not significantly different from zero, the stoma was categorized as <u>SS</u>, if the slope was positive as <u>opening</u> and if the slope was negative as <u>closing</u>. If an upward trend was followed by a downward trend or vice-versa, the stoma was categorized as <u>oscillating</u>.

In the sun period and the final shade period, stomata were categorized as delayed if the lag (λ) in minutes was larger than 0 and the lag fit error was less than 50% of the fitted value.

Stomata in the final shade period were categorized as <u>closing to a QSS</u> if the final QSS fit value was reached with confidence and further data trend did not fall outside fit error for the QSS value. If instead they did not reach a QSS, they were categorized as <u>closing</u>. If further data trend fell outside the fit error for the QSS value and were higher than the QSS, they were categorized as <u>closing and re-opening</u>.

#### Maize

Stomata were categorized as oscillating between open and closed if they fully closed and re-opened or opened and fully closed in the initial growth light period (I) (e.g. in SM Fig. 1B).

Stomata were categorized as opening and oscillating (saturating light, period II) or closing and oscillating (final growth light, period III) if after initial opening/closing their apertures were stagnating and thereafter opening/closing further for more than five time-points in a row.

### Morphological measurements

Guard cell length (GCL) was measured in Fiji by manual use of the straight-line tool in the image at the last timepoint of the period with 1000 (Chrysanthemum) or 1500 µmol photons m^−2^s^−1^ (Maize), the timepoint where stomatal aperture was generally maximal. GCL was taken as a proxy for stomata size because it was clear for all stomata. In contrast, guard cell width and surface area suffered from the unclear borders of guard and pavement cells.

A Voronoi plot (Fig. 3E), which connects lines with equal distance to the borders of each neighbouring stoma, was created via the Voronoi tool (in the Fiji software), and each surface was measured via the ‘analyse particles’ menu. Resulting data were used to test for relationships between the leaf area that could be assigned to a given stoma (‘Voronoi area’) and that stoma’s aperture and kinetics. Only Voronoi surface areas that were fully inside the image were considered in this analysis.

Because our method depends on the automated focus projection of the z-stack (the reduction into a single image that represents the entire surface within the FOV), we compared it to the human operator: manual selection of the best focus position per pore. We compared the results for ten stomata with a good distribution within the FOV for 100 stacked images per 119 time points during shade-sun-shade transition for *Chrysanthemum*. Further image processing, segmentation and quantification of the pore area were automated and following the identical protocol for both manual selection and focus projection.

### Plant material and growing conditions

#### Chrysanthemum

Experiments were conducted in a previously described growth chamber (Zhang *et al*., 2024). Chrysanthemum (Deliflor Chrysanten, Maasdijk, the Netherlands) plants that had been grown in plastic pots (diameter 14 cm, filled in with potting soil) were cut back at the third or fourth node to allow for the formation of new axillary buds.

#### Maize

Seeds (Tasty sweet F1, Buzzy) were germinated and grown in plastic pots (diameter 14 cm, filled in with potting soil) for one week at 160 *µ*mol m^−2^ s^−1^ followed by two weeks at 500 *µ*mol m^−2^ s^−1^. The photoperiod was 18:6 (light/dark) with temperature 22/20 °C (day/night). Plants were irrigated twice per day (at 9:00 and 17:00 h) with nutrient solution, using an automatic ebb and flow system. Experiments were conducted in a different room because the growth chamber had a layout with stacked layers for plant growth with too little space to fit the microscope. Temperature in the measurement room was 23°C and RH was 70%. Plants were acclimated for at least 60 min in the measurement room before measurements were started.

### Stomatal aperture measurements

Stomatal apertures were measured on the youngest fully expanded leaf (Chrysanthemum) or the oldest developing leaf (Maize) in the middle between the leaf edge and the midrib. A screen was used to shade the measured plant from direct growth chamber lighting during aperture measurements (Chrysanthemum). All lighting and imaging protocols used are reported in the SM Methods section.

### Statistical tests

Normality tests for the distribution of fit or measured parameters were performed with Shapiro-Wilkins test. KWANOVA and paired Wilcoxon signed rank tests were used to test for significant differences between sets of parameters that were not normally distributed. Spearman’s correlation was calculated between sets of parameters that were not normally distributed. For normally distributed data, ANOVA, students t-test and Pearson correlations were used. All statistical tests were performed in Origin (OriginLab).

## Results

Our portable microscope enabled the imaging of stomatal dynamics of Chrysanthemum and Maize by its selective use of green light, with limited effects on stomatal movement (SM Fig. 2) (Jones *et al*., 2022) for imaging at a low light intensity, while using stepwise changes in light intensity (red/blue spectrum) to trigger changes in stomatal aperture.

### Comparison of stomatal pore area dynamics between automatic and manual selection of best focus per pore

The dynamics of the pore area in the focus projection images (automatic) were well correlated with those in the manually selected best single focus images (Fig. 4A, SM Fig. 3, r >97% P<0.001). However, the pore area tended to be smaller when derived automatically: pores were on average ten µm^2^ smaller in automatic than in manual images (Fig. 4B). To investigate if this difference in pore area arose from spatial dependence of image quality, we correlated the difference against positional coordinates of the stomata. We found that these differences between automatic and manual-derived pore area were neither correlated to the x ( r=-0.16, not significant (NS)), y (r=0.2, NS) position of the pore in the image, nor to the distance of the pore to the image edge (ρ=0.24, NS), nor to the position in the stack of the manual images (r=-0.36, NS). Further, pore opening and closing speeds in the automatic images were similar in the manual images, as indicated by the maximum opening and closing speeds of the fitted curves (Opening: automatic 0.99±0.05, manual 0.93±0.04 µm^2^min^−1^, Closing: automatic 2.1±0.3, manual 1.9±0.3 µm^2^min^−1^, NS). In conclusion, the automatic method can correctly assess the kinetic parameters of the stomata but may underestimate the true pore area.

**Figure 4.**
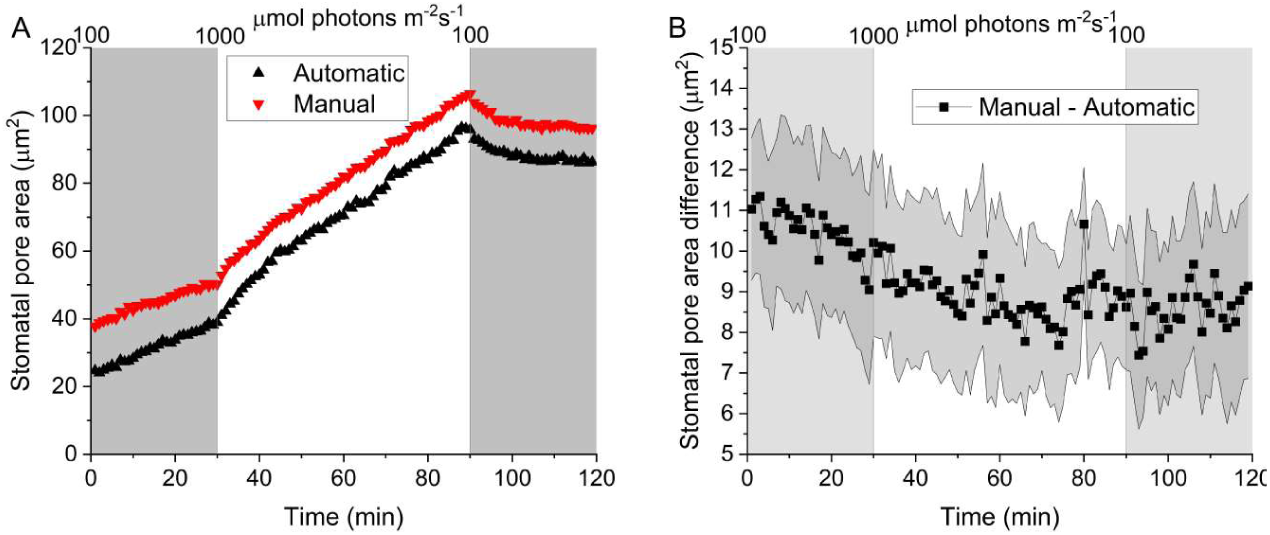
An Example of kinetics of one Chrysanthemum stoma analyzed with the focus projection (automatic) method or in the manually selected best single focus images (manual) demonstrating their correlation. All individual stomata curve pairs are in SM Fig 2. B Averaged difference in pore area between the same pore analyzed in the automatic images and the Manual images in all timepoints of the shade-sun-shade transition. The shaded area around the curve indicates the standard error of means (n=10). Light grey blocks indicate shade periods.

### Movement of the leaf surface upon light intensity change

Because we recorded changes in the position of the leaf surface via automatic adjustments by the step motor of the microscope, we could observe the movement of the leaf surface upon changes in light intensity (Fig. 5). Specifically, after the switch from 100 to 1000 μmol m^−2^ s^−1^ for Chrysanthemum, there appeared to be some shrinkage or bending of the leaf, as its surface moved away from the camera, whereas upon the transition from 1000 to 100 μmol m^−2^ s^−1^, the opposite happened (Fig. 5). These leaf surface movements had no impact on imaging, because the FOV of the leaf was always captured in focus within the stack. Leaf surface movements resulted in a minor displacement of the position of the leaf surface in the stack (SM Fig. 4). Thus, the autofocus software adjusts the z-position of each new image stack based on the previous z-position, because the leaf surface shifts hundreds of micrometers when light intensity changes.

**Figure 5.**
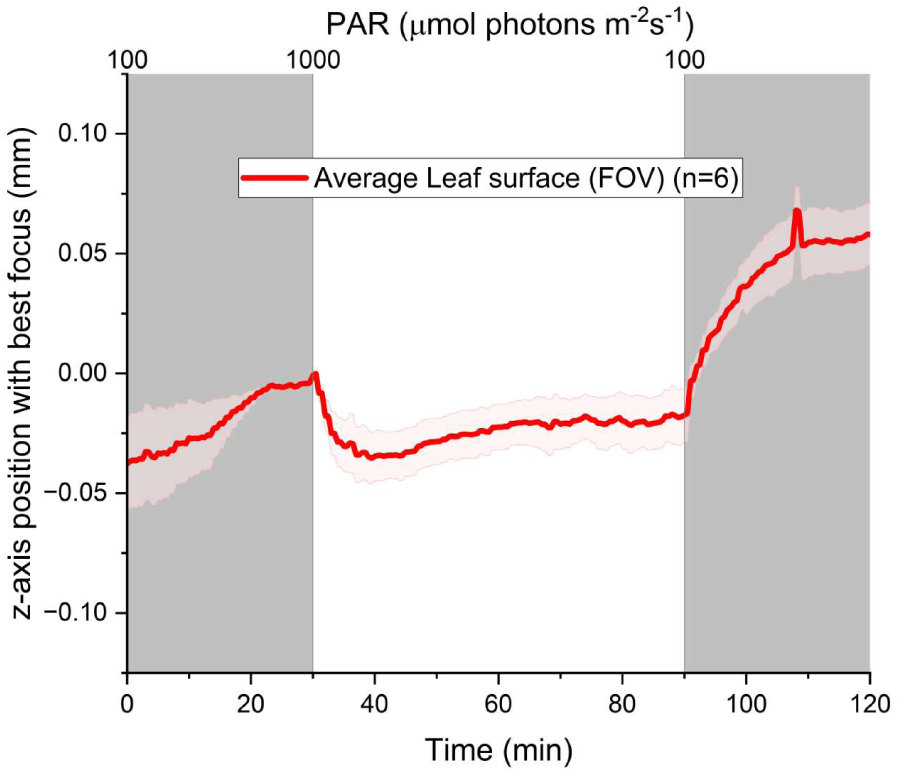
Position on the z-axis of the imaged area of the Chrysanthemum leaf surface where the largest area of leaf surface was in focus during the shade-sun-shade experiment. The red curve indicates the average of six biological replicates, and the shaded area indicates the standard error of the mean. The z-axis position at t=30 min was set to zero for each measurement before averaging. The focus position (y-axis) indicates the middle position of 100 images taken per minute, each of which was spaced one µm apart.

### Stomatal dynamics of maize leaves during transitions between growth and saturating light intensity

A central question in research on stomatal biology is whether morphological parameters of individual stomata such as their size (GCL) and the mesophyll area for which they provide gas exchange (Voronoi area) impact their kinetics. To investigate this, we measured these morphological parameters and quantified the kinetic responses of 29 stomata (45%) of four maize plants during a transition from growth light intensity (500 µmol m^−2^ s^−1^) to photosynthetically saturating light intensity (1500 µmol m^−2^ s^−1^), and back to growth light intensity (e.g. in SM Fig 1). 35 (55%) stomata were unresolved of which eleven were on an image edge, 12 opened but pore area could not be segmented, one clustered smaller stoma did not open and for eleven stomata the pore image quality was so poor that it could not be determined if they opened or not. Nine of these stomata with very poor image quality were near a vein.

We observed a number of different responses among the resolved stomata which we then categorized as <u>closed</u>, <u>open</u>, or <u>oscillating</u> between open and closed, under initial exposure to growth light (I), <u>opening towards a new SS</u>, or <u>opening and oscillating</u>, during the saturating light period (II) and <u>closing towards a new SS</u>, or <u>closing and</u> <u>oscillating</u>, during the final period (III) (Fig. 6A). Stomata were observed to have combinations of all categories of period I-III except open (I) – open and oscillating (II) – closing and oscillating (III). Interestingly, GCL was different between all categories in the initial growth light period (I, Fig. 7A). In contrast, no differences were found between kinetic categories in periods II-III for GCL and in periods I-III for Voronoi area (SM Fig 5 A-E). For these stomata, their initial state in the growth light was thus strongly linked to their GCL, but the occurrence of oscillations in their subsequent kinetics was not.

**Figure 6.**
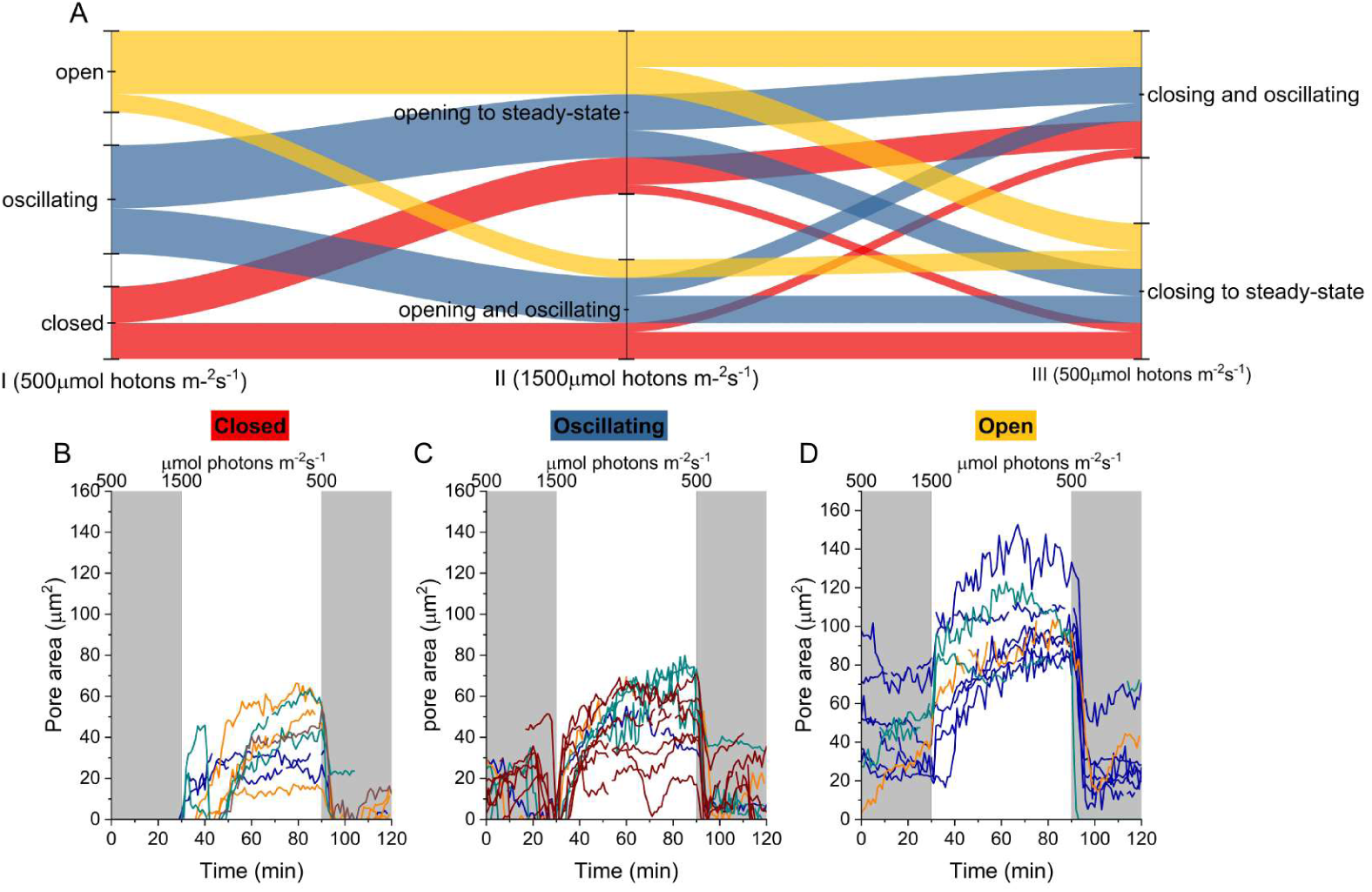
Categorized kinetics of 29 maize stomata during transitions from growth light to saturating light and back to growth light intensity. **A.** Categorized kinetics during the three periods (I) initial growth light 500 µmol photons m^−2^s^−1^ (30 min), (II) saturating light 1500 µmol photons m^−2^s^−1^ (60 min) and (III) return to growth light intensity 500 µmol photons m^−2^s^−1^ (30 min). The thickness of bands in A indicates the fraction of stomata in that category. Colored curves in A indicate how the fraction of stomata within the starting period categories transition to the categories in period II and III. Kinetics in period I were categorized per stoma as being **B** closed, **C** oscillating between open and closed or **D** open. Stomata categories during the saturating light period were opening towards SS or opening and oscillating. Similarly stomatal categories during period III were closing towards SS, or opening and oscillating. Colors of curves in B-D indicate individual plants.

**Fig 7.**
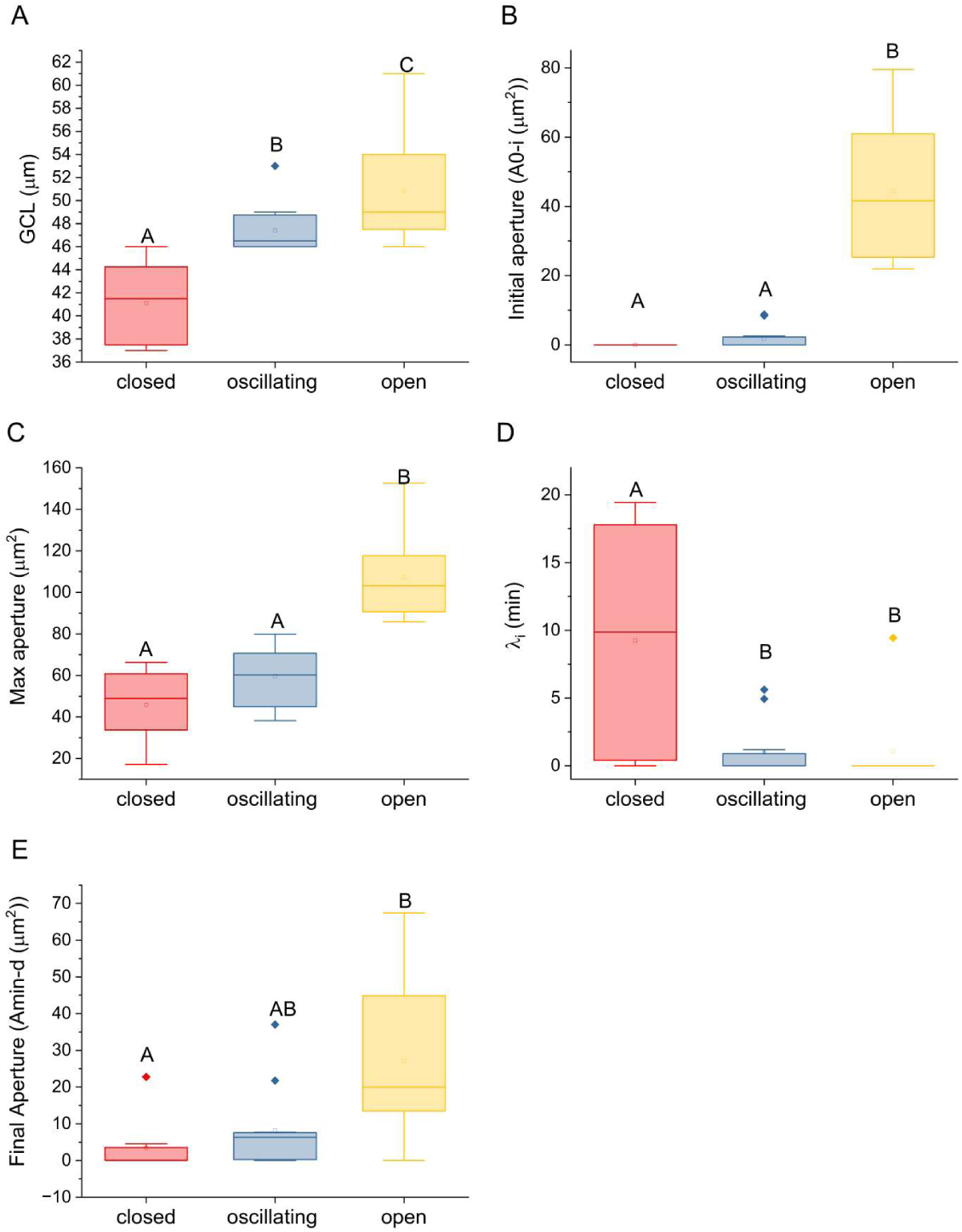
Categories of maize stomatal behavior under growth light intensity, and relationships with morphology and kinetics during light-driven opening and closing of maize stomata. **A** Guard cell length (GCL). **B** Initial pore aperture, **C** maximum pore aperture and **D** lag time (λ_i_) until opening response in the saturating light period (II). **E** pore aperture after initial closing in the final period (III). Significant differences between kinetic categories by ANOVA (A, C) or KWANOVA are indicated by different capital letters (P<0.05). Squares represent the average and lines the median (Q2. Box outlines represent the lower (Q1) and upper quartile (Q3), and the whiskers indicate Q0 and Q4.

The starting aperture and the maximum aperture during the saturating light period (II) were different between stomata that were closed or oscillating and those that were open during the initial period with the growth light (I) (Fig 7BC). In contrast, the lag time before the opening response under saturating light (II) was smaller for initially open or oscillating stomata than for initially closed stomata (Fig 7D). Thus, while both closed and oscillating stomata initially had a low and similar aperture, the oscillating stomata were equally fast responding as the open stomata from the initial period, while the closed stomata were not. No differences were found in aperture or opening lag in saturating light between stomata that were opening with or without oscillations in saturating light (SM Fig 5 F-G).

For period III (1500->500 µmol photons m^2^s^−1^), the apertures after the initial closing response were again minimal for closed and largest for stomata that were categorized as open in period I (Fig 7E). Stomata that were opening or closing with oscillations were not different in aperture after the initial closing than those without oscillations (SM Fig 5 I-K).

### Speed of stomatal movement

Maximum opening speed in period II and closing speed in period III were not different between any of the stomata in the categories of periods I-III (SM Fig 6 A-F) and are therefore presented for all stomata in Fig 8A. Unlike the maximum opening speed, the maximum closing speed was positively correlated to guard cell length (ρ=0.51, p=0.01). The average maximum closing speed was seven times larger than the average maximum opening speed, and maximum opening and closing speeds were positively correlated (ρ=0.47, p=0.03). Similarly, the time constant of stomatal opening, k_i_, was eight times larger than the time constant for stomatal closure, k_d_ (Fig. 8B). Average maximum opening and closing speeds were underestimated, because the fastest stomata (five opening and six closing) were excluded from the speed analyses, as their speed could not be fitted correctly. These stomata opened or closed to a SS within one time point with a large difference in aperture (e.g. k_i_ <0.5 min). To eliminate that these fast responses were not processing errors, we again compared the automatic processing to the manually selected best focus per pore. The images of the time-points at the transition as well as the kinetics (e.g. in SM Fig. 7) validate the absence of processing artefacts. These stomatal responses are thus truly speedy outliers.

**Fig. 8.**
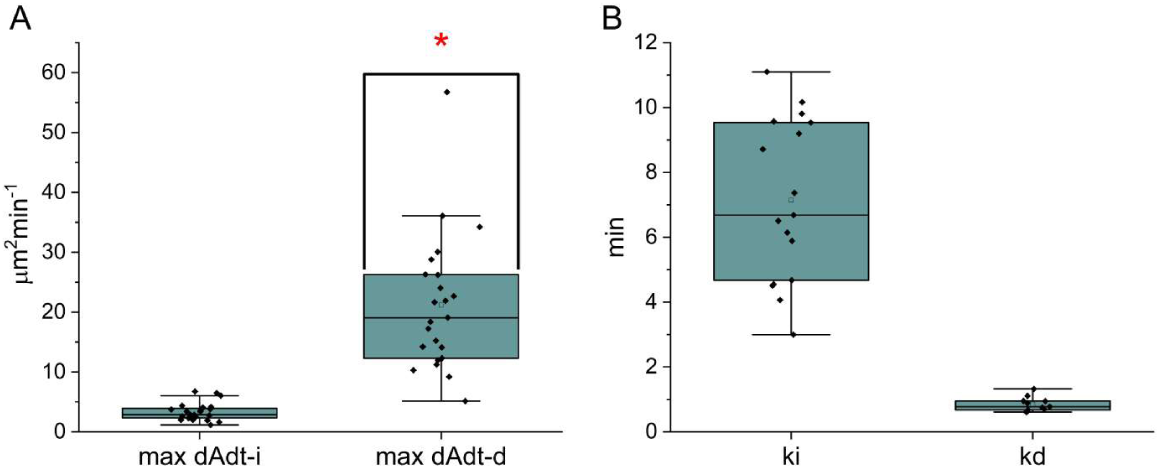
Parameters characterizing rates of stomatal opening and closure in maize stomata during transitions from 500 to 1500 back to 500 µmol photons m^2^s^−1^. **A** maximum opening and maximum closing speeds (n=23). Asterix indicates significant correlation with GCL (ρ=0.51, p=0.01). **B** k_i_, opening time constant for stomata that opened to SS (n= 17). k_d_, closing time constant for stomata that closed to SS (n= 11). Squares represent the average and lines the median (Q2. Box outlines represent the lower (Q1) and upper quartile (Q3), and the whiskers indicate Q0 and Q4.

In conclusion, the annotation of the considerable variation in the kinetics of individual stomata demonstrates the ability of our method to identify and quantify relations between morphological parameters such as guard cell length and kinetics for individual stomata.

### Stomatal dynamics of *Chrysanthemum* leaves during a shade-sun-shade transition

To further demonstrate the power of our method, we tested it in a shade-sun-shade transition on leaves of Chrysanthemum, where photosynthesis typically reaches a SS during the sun phase (60 min), but stomatal aperture does not (SM Fig 2 (Zhang *et al*., 2022, 2024)). The rate of stomatal opening at 60 min of high light is higher and in absolute terms more variable than close to SS pore area at 180 min for individual stomata (SM Fig. 8, (t=60 min) 1.25 ± 0.09 > (t=180 min) 0.25 ± 0.05 µm^2^min^−1^). Therefore, we investigated the effect of the change in irradiance on individual stomata, when individual apertures and rates of opening were more heterogeneous than at steady state, to derive relations to subsequent responses during the shade phase. We imaged the leaves of six plants split over two cultivation periods. After image processing, 124 opening and 95 closing kinetics out of 158 stomata in the FOV were resolved (Fig. 9).

**Figure 9.**
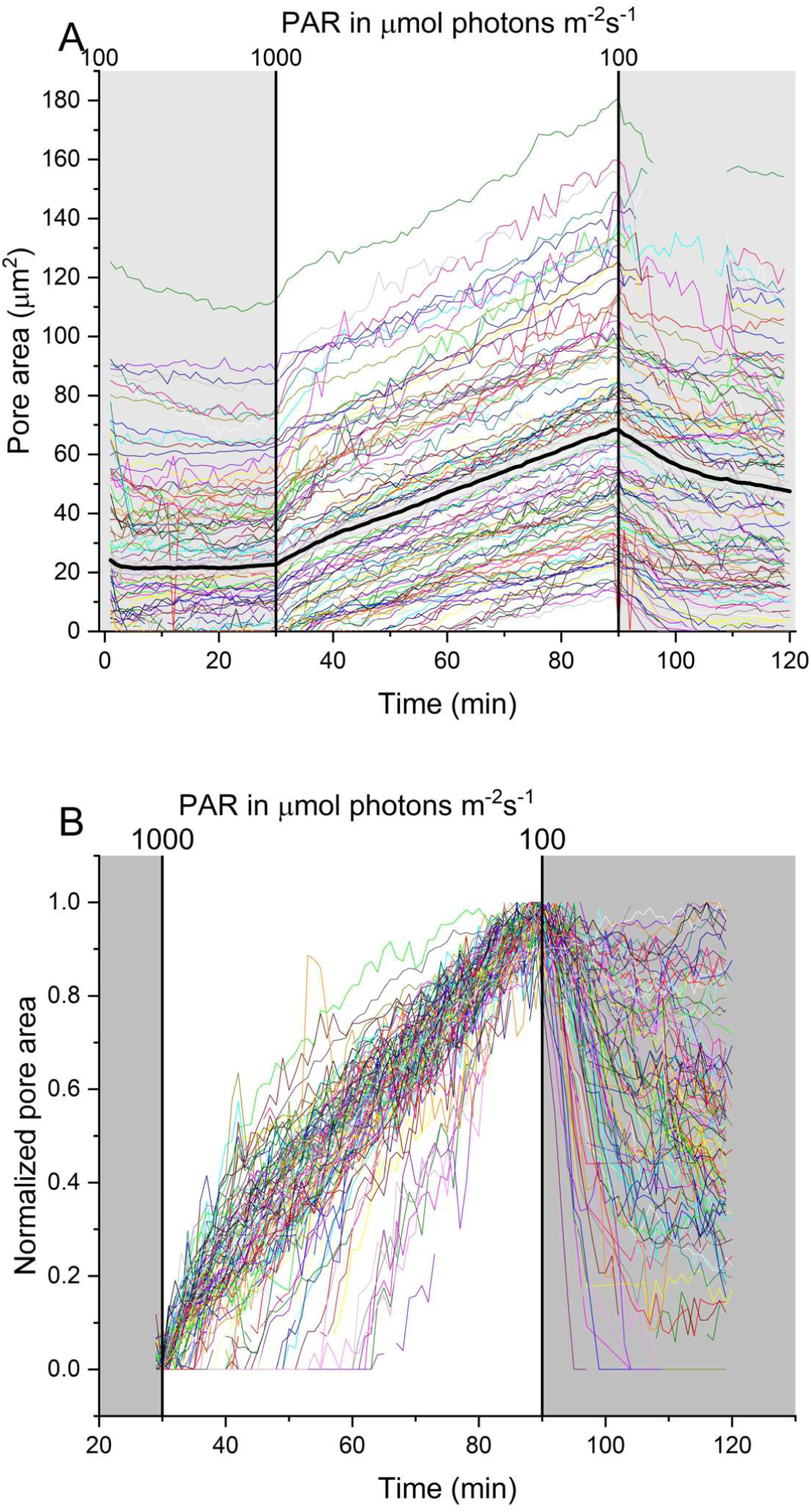
Stomatal pore area changes upon a shade-sun-shade transition. Single lines show values of 124 individual stomatal pores from six chrysanthemum plants. The thick black curve represents the average, and the grey area the standard error of the mean. During the second shade period, a temporary loss of focus during one measurement caused fewer data points to be recorded. B Kinetics of all stomata shown in A, normalized to 0 at t=29 min just before the light intensity was increased and to 1 for the maximum aperture reached at 1000 µmol photons m^−2^s^−1^, highlighting the differences in opening lag (λ_i_) and stomatal closure.

While average stomatal aperture was in a SS in the initial shade condition (Figure 9, 0-30 min), some individual stomata showed minor movements by either slightly opening or slightly closing. 47 stomata were closed, 25 were open in a SS, 27 were opening, 21 were closing and one stoma was open but oscillating. Closed stomata had smaller GCL than those that were closing in initial shade (Fig. 10A). In addition, closed stomata also had smaller Voronoi area than those that were closing or in a SS in initial shade (Fig. 10B). Apertures at the start of the sun period were not different between opening, SS, closing or oscillating stomata in the initial shade (SM Fig. 9A).

**Fig. 10.**
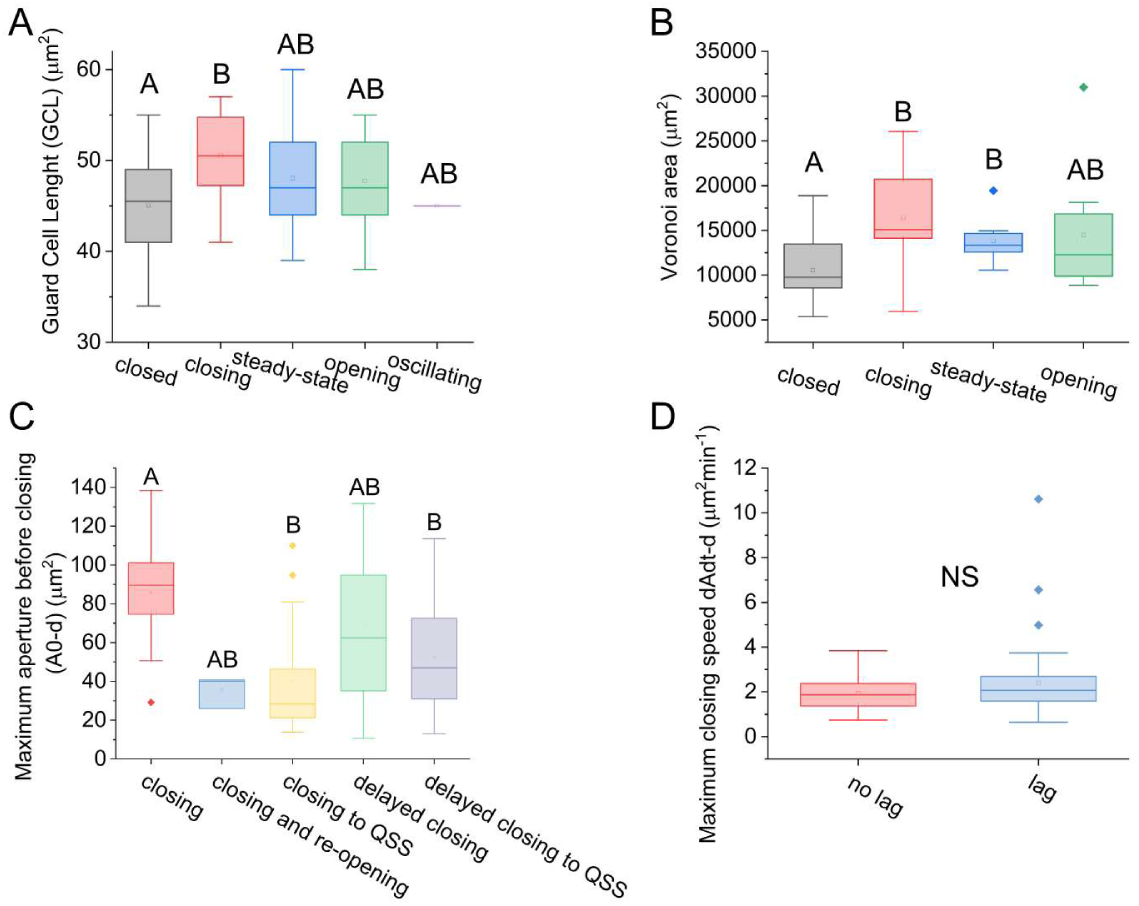
Morphological and kinetic parameters of categorized Chrysanthemum stomata during a shade-sun-shade transition. **A** Guard Cell length (GCL) and **B** Voronoi area compared between the kinetic categories of the initial shade phase. Number of stomata per category: 47 closed, 25 open in a steady-state, 27 opening, 21 closing and one stoma was open but oscillating. Voronoi area could only be quantified for 27 closed, 12 open in a steady-state, ten opening and 9 closing stomata **C** Maximum pore aperture in the sun phase before closing. Number of stomata per category: 12 closing, three closing and reopening, 16 closing to QSS, 15 delayed closing and 38 stomata in delayed closing to QSS. **D** Maximum pore closing speed compared between categories with and without closing lag (λ_d_) of the final shade phase. Squares represent the average and lines the median (Q2. Box outlines represent the lower (Q1) and upper quartile (Q3), and the whiskers indicate Q0 and Q4.

After the light intensity was increased, 70% of resolved stomata (87) immediately opened their pores towards a SS, while the rest showed a lag in their response of up to 40 min (Fig. 9). Most stomata with an opening lag were closed in the initial shade (SM Fig 9B). Maximum opening speeds between stomata with and without delayed opening were not different (SM Fig 9C). Maximum aperture in the sun phase was lower for initially closed stomata than for opening, closing or SS stomata in the initial shade (SM Fig. 9D). Similarly, stomata with a delayed opening response reached a lower maximum aperture during the sun phase (SM Fig 9E).

The light intensity decrease led to an immediate closing response of 32 stomata (33%), of which 18 reached a quasi-steady-state (QSS), four closed and re-opened and ten did not reach a QSS (all ten were on the same leaf). We use QSS here, because the average final shade aperture was larger than the average initial shade aperture (A_min-d_ 23±3 µm^2^ > A_0-i_ 13±2 µm^2^, mean+SE, paired sample Wilcoxon Signed Ranks Test, P<0.005), and because further closing could have occurred after 30 min in the final shade. Immediately closing stomata that did not reach a QSS had a larger aperture at the time of the light intensity decrease than those that reached a QSS (Fig 10C).

53 stomata (54%) had a delayed closing response, of which 38 reached a QSS with no differences in aperture at the time of the light intensity decrease between these two groups (Fig. 10C). No indication was found that delayed stomata could compensate for their delay by a faster closure, because the maximum closing speed was similar between delayed and immediately closing stomata (Fig. 10D). 19 stomata (19%) did not close at all within the 30 min sun-shade transition. These stomata were not evenly distributed among the biological replicates (numbers per replicate: 15:1:3:0:0:0) and had a similar aperture at the time of the light intensity decrease than the stomata in the other closing categories (74 ± 5 µm^2^).

Of all parameters, only guard cell length was normally distributed (Shapiro Wilkins test, p<0.001), therefore Spearman’s nonparametric ρ was used to investigate correlations. A larger aperture during the first shade phase (A_0-i_) was strongly correlated with a larger maximum aperture in the sun phase A_0-d_, and moderately correlated to the QSS aperture during the second shade phase A_min-d_ (Fig. 11). A_0-d_ was also moderately correlated to the maximum opening speed, dAdt_I_ (Fig. 11). The time lag of the opening response (λ_i_) was negatively correlated to A_0-I_, GCL, Voronoi, A_0-d_, A_min-d_, λ_d,_ and positively to the maximum closing speed, dAdt_d_ (Fig. 11). Interestingly, the time lag of the closing response (λ_d_) was correlated to A_0-i_ and A_0-d_ (Fig. 11).

**Figure 11.**
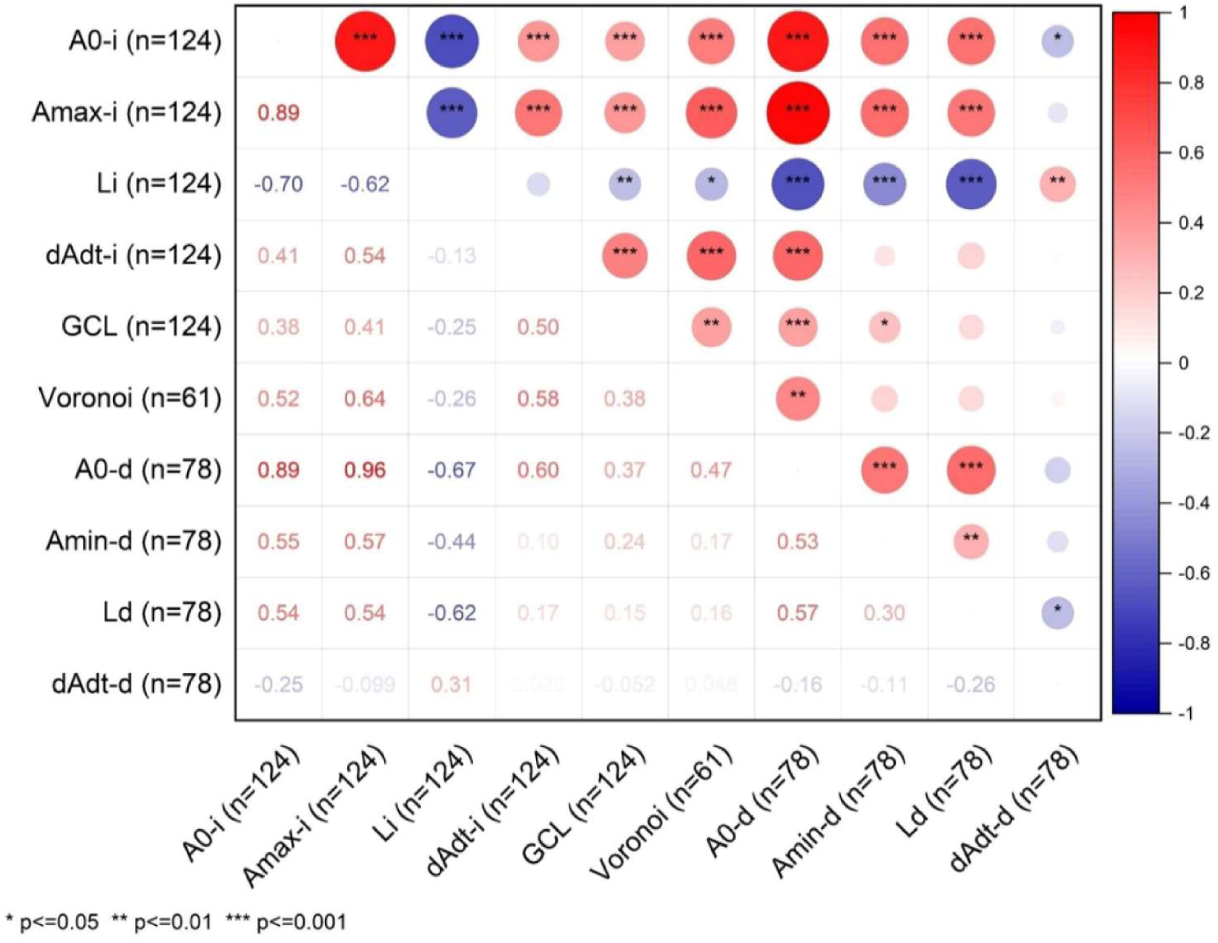
Spearman’s correlation matrix between parameters of individual Chrysanthemum stomata (based on Fig. 9). A_0_, fitted steady-state aperture under 100 µmol photons m^−2^s^−1^ PPFD. A_max_, maximum aperture under 1000 µmol photons m^−2^s^−1^ PPFD; Li/Ld, lag (λ_i_/ λ_d_) in the response time of stomatal aperture to a step increase or decrease in PPFD; dAdt, maximum rate of stomatal opening (-i) or closing (-d) to an increase or decrease in PPFD from 100 to 1000 µmol photons m^−2^s^−1^ or vice versa. Anatomical parameters of guard cell length (GCL) and Voronoi area were also compared. The Voronoi area represents the fraction of the leaf surface that is within the shortest distance to that stoma. A_0-d_, fitted final aperture under 1000 µmol photons m^−2^s^−1^, A_min-d_, predicted final steady-state aperture under 100 µmol photons m^−2^s^−1^ (QSS).

Among the anatomical parameters, Guard cell length (GCL) was moderately correlated to the maximum opening speed dAdt_max-i_, initial A_0-I_, maximum A_0-d_, and final aperture A_min-d_, as well as the Voronoi area (Fig. 11). Additionally, the Voronoi area was moderately correlated with A_0-i_, A_min-d_ and the maximum opening speed dAdt_max-i_, (Fig. 11).

In conclusion, our method resolved ten different kinetic and non-kinetic parameters for up to 78% of the *Chrysanthemum* stomata within the FOV to generate new insights into the WUE of individual stomata.

## Discussion

We have presented a real-time *in planta* microscope, which provides a new way to study the opening and closure of individual stomata in the growth environment. This method facilitates analyses of individual stomatal dynamics in relation to their local morphology (Voronoi) and anatomy (Guard Cell Length). Our hardware and method are distinct from the few previous studies that have imaged stomata *in planta* (Kaiser and Kappen, 1997, 2001; Kaiser, 2009; Grantz, Zinsmeister and Burkhardt, 2018). The main differences to previous approaches, as well as their advantages and disadvantages, are listed in Table 1.

**Table 1.** Differences between this work and prior studies (Kaiser and Kappen, 1997, 2001; Kaiser, 2009; Grantz, Zinsmeister and Burkhardt, 2018) with their advantages and disadvantages.

| Difference | Advantage | Disadvantage | Study |
| --- | --- | --- | --- |
| **525 nm** versus 880 nm imaging light | Higher optical resolution | Imaging light is photosynthetically active, closing effect on stomata | (Kaiser & Kappen, 1997, 2001) |
| **Autofocus per stack** instead of per pore | Imaging speed is not limited by number of stomata, but by stack size | Can cover stomata in only a small area of the leaf (~1 mm<sup>2</sup>) | (Kaiser & Kappen, 1997, 2001; Kaiser, 2009; Grantz et al., 2018) |
| **Kinetic model fitting** | Quantitative comparison of fit parameters | - |  |
| **No Gas exchange chamber** | Boundary layer thickness is close to natural | No information on leaf bulk gas exchange | (Kaiser & Kappen, 1997, 2001; Kaiser, 2009; Grantz et al., 2018) |

A major contrast with prior studies is that the way the leaf is clamped was minimally invasive to the microenvironment of the leaf, with free transpiration on both sides of the leaf, because no gas exchange measurements were performed. Unlike in gas exchange cuvettes that use fans to stir the air and thus minimize boundary layer thickness, in our method the thickness of boundary layer was not regulated, reflecting the natural state of the leaf. Another major difference was the sampling of stomata: prior studies have randomly sampled stomata over a large area (>1 cm^2^), whereas we resolved as many stomatal kinetics as possible within ~1 mm^2^, to relate their kinetic behavior to the local morphology on the surface of the leaf (e.g. Voronoi area). We reached a higher time resolution of, on average, 21 stomata min^−1^ for *Chrysanthemum*, but seven stomata min^−1^ for maize, compared to 75 stomata once every 6-7 min in *Vicia Faba* (Grantz, Zinsmeister and Burkhardt, 2018). Sampling in ~1 mm^2^ means however that variations in dynamics over greater distances are missed. Another major difference to previous studies is that we fit a kinetic model to individual stomatal aperture kinetics, to be able to quantitatively compare them and derive functional relationships. Further, our method detects changes in leaf thickness or bending under dynamically changing light intensity (Fig. 5), a feature that may enable it to be used in studies investigating functional relationships between leaf morphology, leaf water relations, and stomatal pore area (including dynamic changes in all of these properties).

Our method measured smaller pore area compared to manual selection of the best focused single image per pore (Fig. 3). The smallest part of the pore may, however, not necessarily be entirely in focus in a single image, especially for large stomata. It could therefore be plausible that the true pore area lies between both types of measurements of pore area. Implementation of the state-of-the art in computer vision solutions, such as StomaAI (Sai *et al*., 2023), for stomatal measurements and segmentation may increase the resolving power.

### Comparing Maize and Chrysanthemum

In contrast to Chrysanthemum, where 78% of stomata were clearly resolved, a significant portion of stomata in maize (55%) remained unresolved. This discrepancy was partly due to the more transparent leaf surface in maize, which allowed structures beneath the surface to be visible. As a result, the surface projection algorithm faced challenges, producing lower-quality images compared to those of Chrysanthemum, where this issue did not arise.

The GCL for Maize and Chrysanthemum was on average 47 µm (Fig. 7 A, Fig. 10A), with either dumbbell (Maize) or elliptic (Chrysanthemum) anatomy. Although their length was similar, the kinetics were much faster in Maize stomata compared to Chrysanthemum, with higher opening and closing speeds and less lag (Fig. 9, SM Fig 8 C, D), related to their anatomy (Lawson and Vialet-Chabrand, 2019). In addition, the different light intensity changes and the different leaf age and developmental stages may have affected the relative difference in kinetics.

Stomatal size was found to be negatively correlated with WUE across different Arabidopsis ecotypes (Dittberner *et al*., 2018). A high WUE is typically associated with a fast rate of stomatal closure (Lawson and Vialet-Chabrand, 2019), and smaller stomata have indeed often been observed to open or close faster than larger stomata (Drake, Froend and Franks, 2013; Kardiman and Ræbild, 2018; Durand *et al*., 2019). In contrast, we found a strong positive correlation between GCL and maximum opening speed for Chrysanthemum as observed before for elliptical/kidney-shaped stomata (McAusland *et al*., 2016), as well as a positive correlation between GCL and maximum closing speed for Maize. Maximum opening and closing speeds of individual stomata were correlated in Maize (r=0.48, P<0.05) but not in Chrysanthemum (Fig. 11). The relation between stomatal size and speed may thus be more complex than previously assumed. This relation could be species dependent, as no correlation between stomatal size and speed was found in a diverse range of plants with differing stomatal morphologies and physiological behaviors (Haworth *et al*., 2018). Additionally, it may depend on the type of stimulus, since e.g. step changes in light intensity and in VPD often trigger different dynamics (Durand *et al*., 2019).

Average Voronoi area was smaller for Chrysanthemum stomata (13000 µm^2^, Fig. 10B) compared to Maize stomata (28000 µm^2^, SM Fig. 5D). Similarly, stomatal density was higher in Chrysanthemum images (72 mm^−2^) than in Maize (32 mm^−2^).

### Could stomatal pore oscillations in low light be a means to increase the response speed of stomata to a high light stimulus?

Maize stomata that oscillated between open and closed state in low light intensity and were closed before the light intensity changed responded faster to a high light intensity than stomata that were initially closed in a steady-state (Fig. 7 B, D). The faster opening could have benefits for carbon assimilation (Lawson and Blatt, 2014; Lawson and Vialet-Chabrand, 2019). Oscillations originate from the gain in feedback regulation (Kaiser and Paoletti, 2014; Peak, 2023); they are the result of an interaction of hydraulic effects and active osmotic adjustment of guard and epidermal cells. Oscillations have been argued to allow for a faster response to environmental fluctuations (Kaiser and Paoletti, 2014), and our data seems to confirm this for a response to high light (Fig. 7). It would be interesting to explore this phenomenon further, e.g. by using larger numbers of stomata and stimuli.

### Closing lag and partial closing affect water loss of Chrysanthemum stomata under dynamic light intensity

The analysis of dozens of *Chrysanthemum* stomata produced several insights of stomatal behavior, with important repercussions for WUE: removing the average closing delay of 4.4 min would lead to 13.5% water saving during that 30 min period per average stoma, with a delayed closing response (Fig. 12). Moreover, removal of the partial closing to a QSS would save ~20%, and in combination with removal of the closing lag it would save ~40%, assuming that the maximum speed was kept the same (Fig. 12b). The extent of water saving depends on QSS relative to SS in the shade, and on the closing speed. Considering that delayed closing (51%) and partial closing to a QSS (52%) occurred frequently, it would be interesting to find out if these targets could really be used to improve WUE (in Chrysanthemum and elsewhere). Moreover, larger aperture in high light intensity (A_max-i_) correlated positively with the time lag for the closing response (λ_d_), further enhancing the consequence for WUE. Finally, closing delays were longer for Chrysanthemum stomata that were close to the SS pore aperture at 1000 µmol photons m^−2^s^−1^ (SM Fig. 8) than for those that were not (Fig. 9) (λ_d_ (180 min ‘Sun’) 14±1 >> (60 min ‘Sun’) 4.4±0.4 min, P<0.0001, KWANOVA) with much larger apertures ( (180 min ‘Sun’) 202±7 >> (60 min ‘Sun’) 56±4 µm^2^, P<0.0001, KWANOVA) and therefore a relatively larger effect of closing delay on WUE at the SS in the ‘Sun’. The larger closing delay for stomata near SS than far from SS at 1000 µmol photons m^−2^s^−1^ indicates that the closing delays were not merely related to hysteresis, e.g. switching from a fast rate of opening to fast rate of closing, but to the absolute stomatal aperture.

**Fig. 12.**
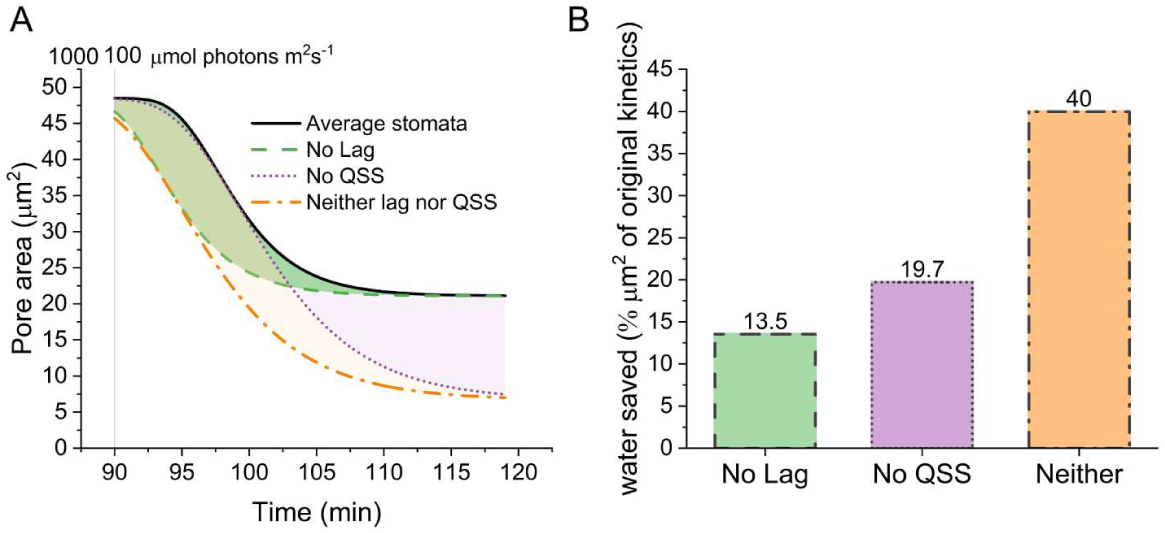
Simulated impact of closing lag and partial closing of average Chrysanthemum stomata on water loss. **A** Modeled closing response with Eqn. 2 of the average stomata with delayed partial closing to the QSS. Average values were taken for maximum aperture in high light, average QSS, average time-constant for partial closing (k_d_) and the average closing lag λ_d_. λ_d_ was set to 0 for no lag without changing other parameters. For no QSS, QSS was changed to the average SS (the initial average aperture in low light) and the average time-constant was adjusted to maintain the same maximum closing speed. For neither lag nor QSS, parameter changes made for no lag and no QSS were performed simultaneously. **B** Color coded areas between the curves in A normalized to the area below the average curve indicate the water saved compared to the average stomata

Across a number of species, Deans *et al* (Deans *et al*., 2019) found that a large stomatal conductance at high light intensity was correlated with fast closing response, to compensate in water-use efficiency. We could not corroborate this for the aperture before closing (A_0-d_) and their maximum rate of closing (max dAdt_-d_ in individual Chrysanthemum stomata (Fig. 11), but we could for Maize (r=0.44, P=0.03). Chrysanthemum stomata with a large aperture thus have a disproportionately large negative contribution on WUE normalized for their apertures, due to the observed correlation between the aperture before closure (A_0-d_) and the time lag for the closing response (λ_i_), combined with a similar closing speed as stomata with a smaller aperture (Fig. 10D).

## Conclusions

We showed here that *in planta* microscopy of stomata could be used to generate stomatal opening and closure dynamics of dozens of neighboring individual stomata. Our method, which uses green light for imaging and generates focus projections of the entire leaf surface within the FOV, resolved the kinetics of 78% of *Chrysanthemum* and 45% of Maize stomata, on average 21 and 7 stomata min^−1^ during light intensity transitions. We detected bending of the leaf and/or changes in leaf thickness, likely related to leaf turgor. Pore area kinetics were fitted with a model developed for stomatal conductance time courses and show the substantial variation between individual stomata. Oscillations of maize stomata were found to contribute to their faster response upon light intensity change. Chrysanthemum pores with larger apertures contributed disproportionately to a lower WUE due to their larger lag before closing than for stomata with smaller apertures. Closing lag and partial closure in Chrysanthemum even contributed on average to a 40% higher aperture area in the first 30 minutes of closure than without these features, wasting considerable water.

## Acknowledgements

We thank Seven-steps-to-heaven for their in-kind support of the actinic light source. We acknowledge 4TU HTSF Plantenna for financial support. Silvere Vialet-Chabrand and Ep Heuvelink are acknowledged for helpful discussion of the results.

## Author contributions

Conceptualization T.B. and J. S., Hardware T.B. R.S. Software T.B., R.S., Growing facilities E.K., Experimentation T.B., Data Analyses T.B., Writing the first draft T.B., Critical review of the manuscript T.B., E.K., J.S., Funding J.S.

## Data availability

All data is available on request. For more information and software updates: GitHub - Plantenna/Stomata-microscope: Non-invasive microscopic imaging of individual stomatal kinetics in the growth environment with high resolution

## Supporting Information

### SM Methods Analyses

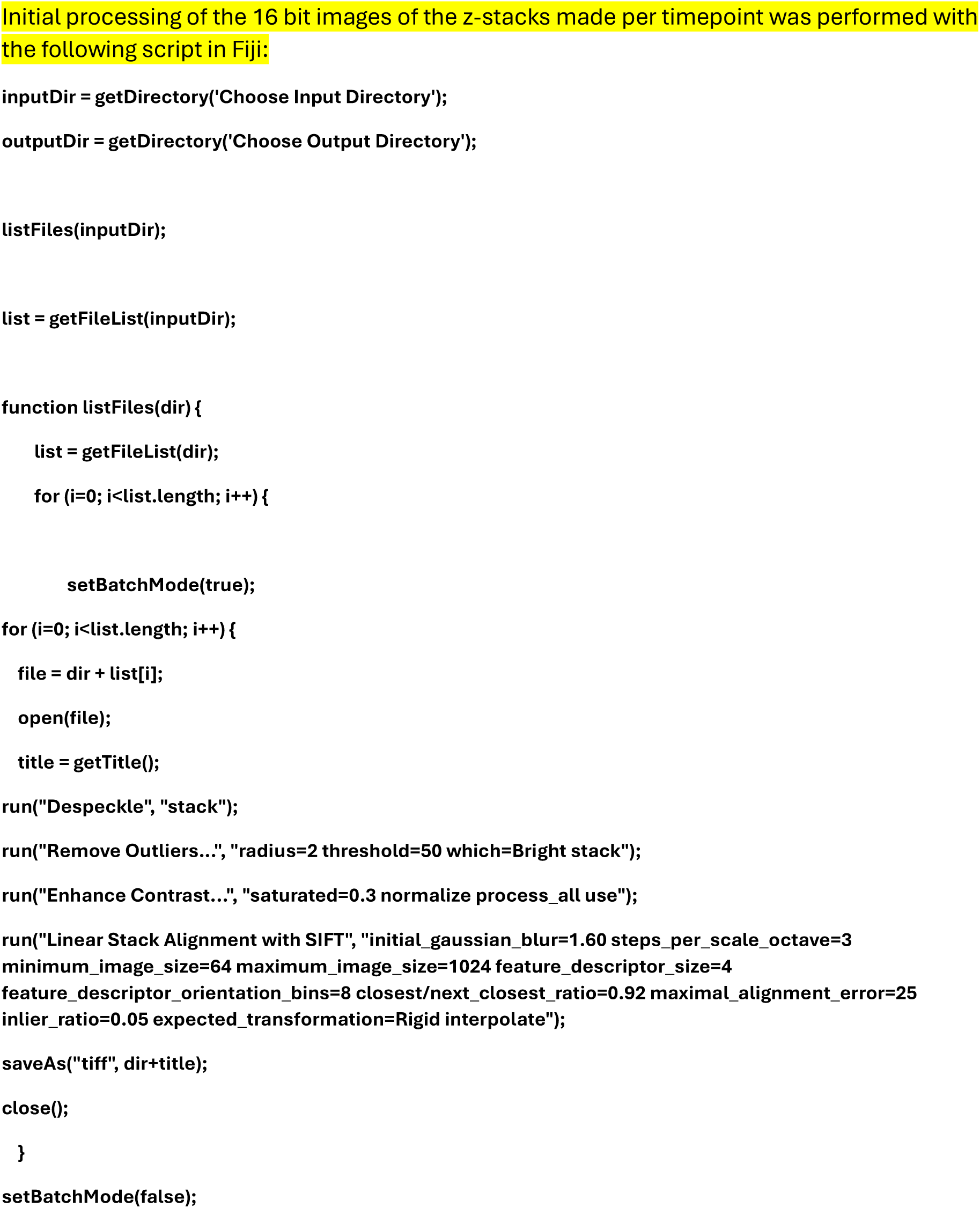
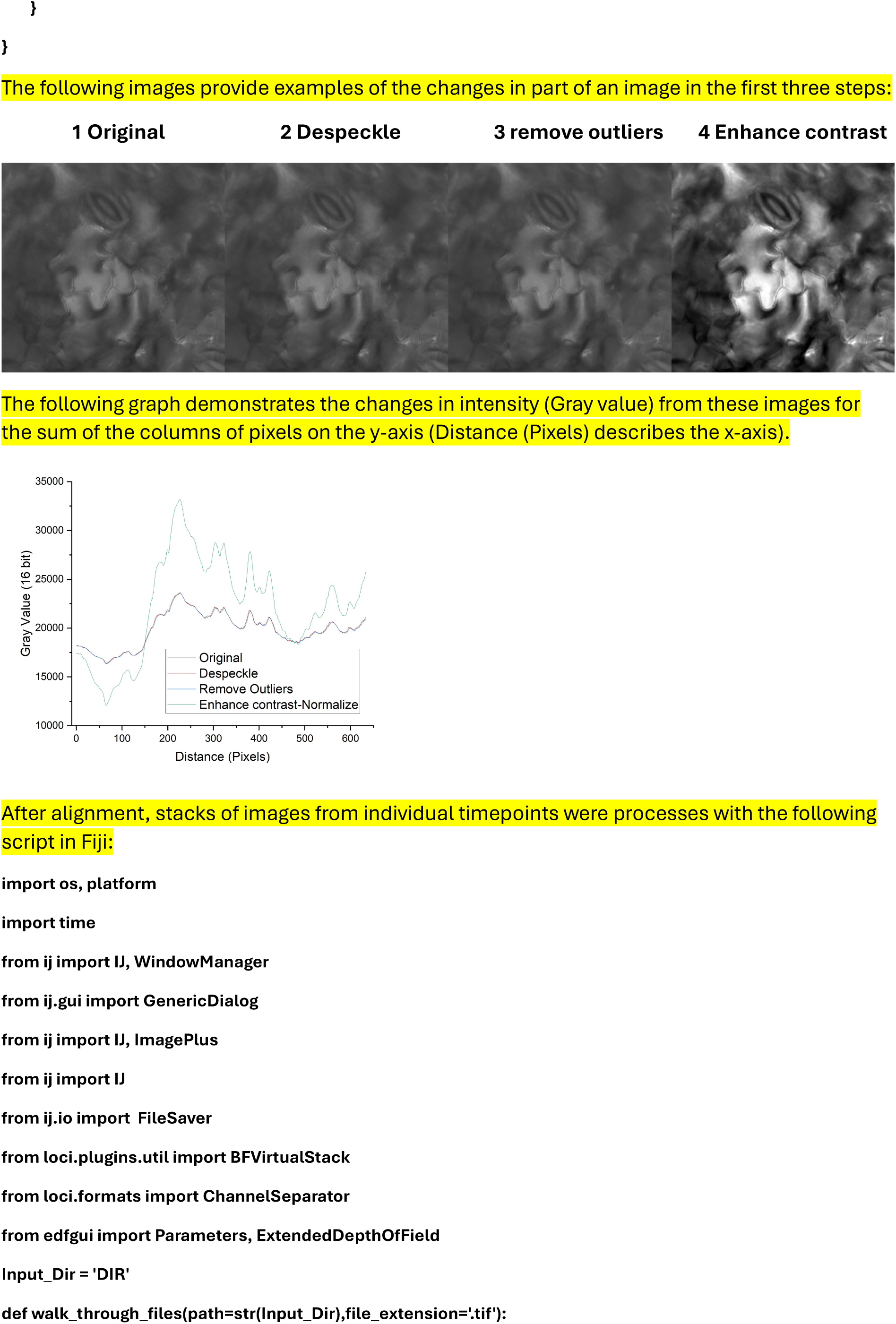
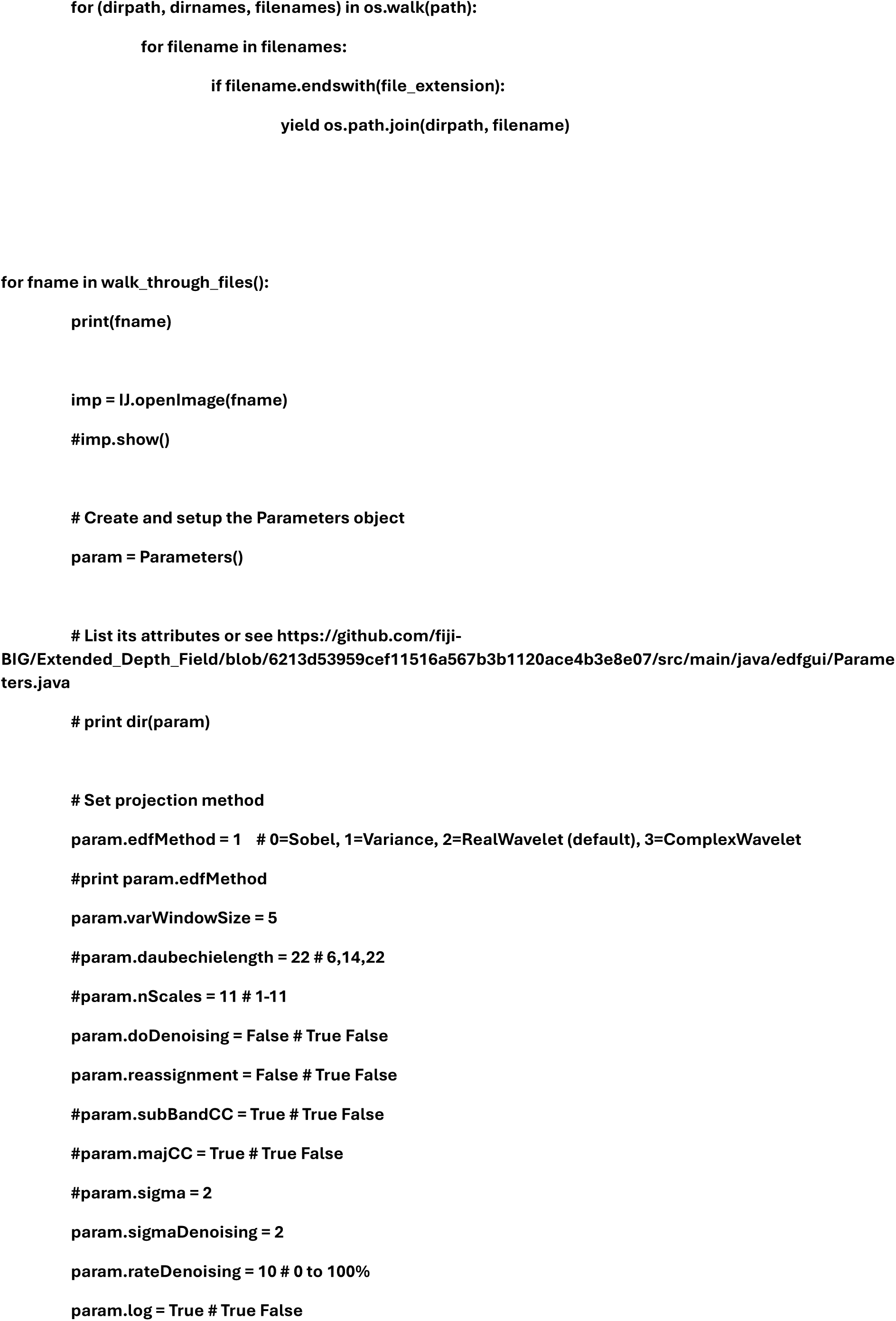
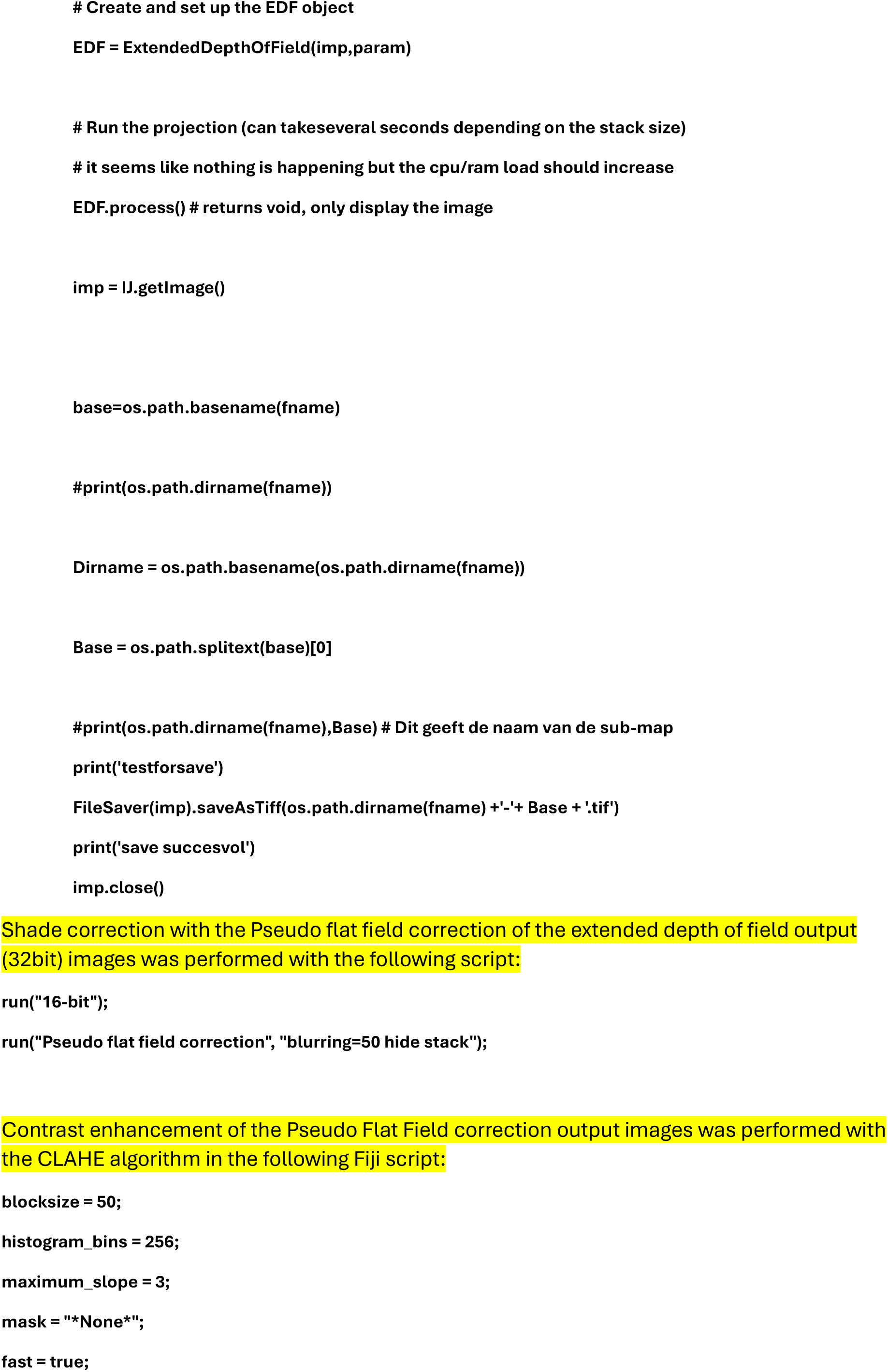
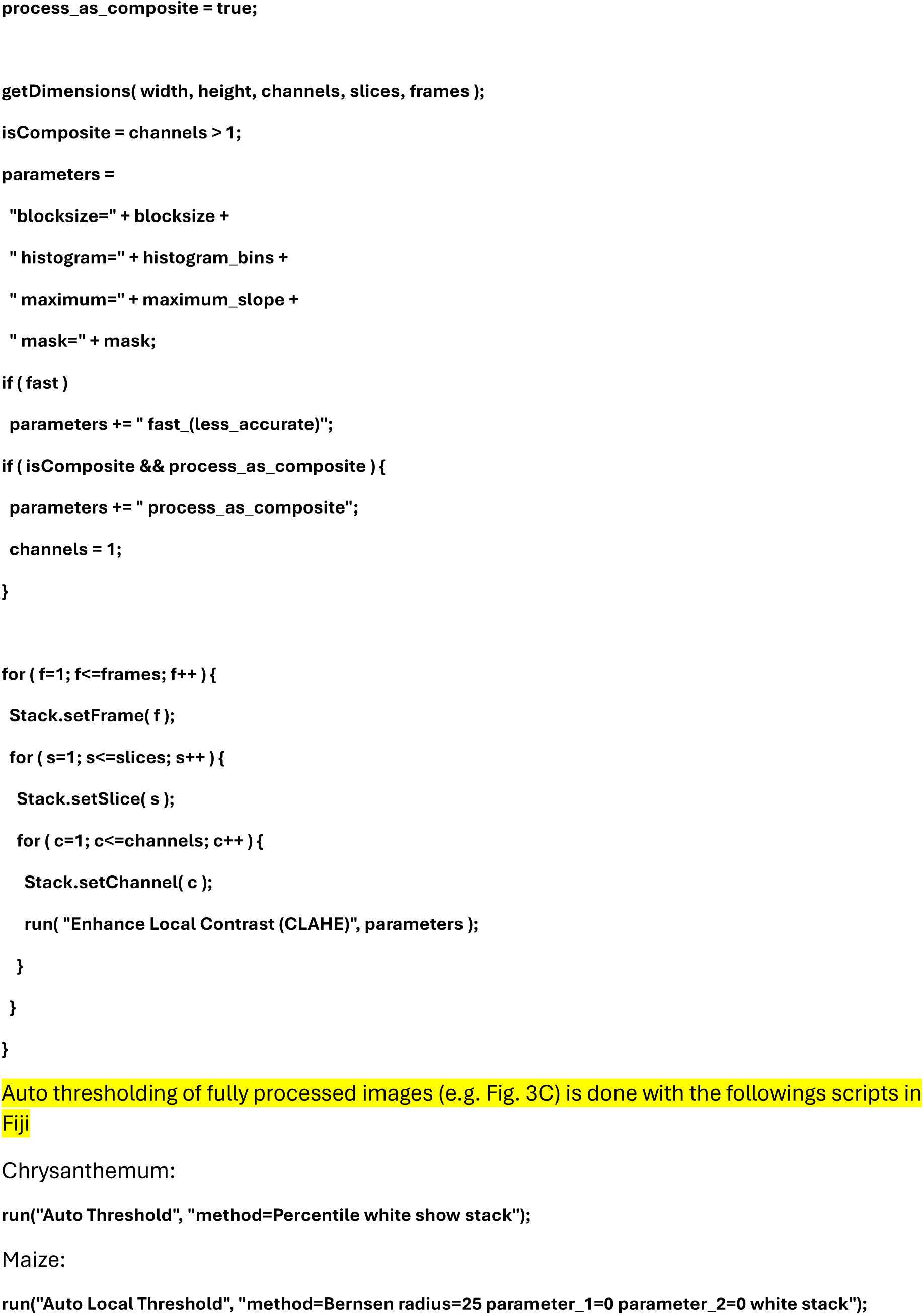

### Lighting and imaging protocols

To fully characterize photosynthetic induction, it is important to reach a steady-state in photosynthesis and stomatal conductance at a given light intensity before switching to a higher light intensity. However, individual stomata often open or close when bulk stomatal conductance is in a steady state. Because we could not measure gas exchange simultaneously with aperture, we were unable to determine whether stomatal conductance was in a steady state. Therefore, we took the times to steady state (see below) from literature ( McAusland et al., 2016; Zhang et al., 2022).

#### *Chrysanthemum* (shade-sun-shade, 180 min)

Leaves were acclimated to 50 μmol m^−2^ s^−1^ blue-red light and 50 μmol m^−2^ s^−1^ green imaging light for 60 minutes before image acquisition was started. Image stacks were acquired every minute for 120 minutes. After 30 (90 total) minutes, the light intensity was increased to 1000 μmol m^−2^ s^−1^ by increasing the blue-red light intensity to 950 μmol m^−2^ s^−1^. After another 60 (150 total) minutes the blue-red light intensity was switched back to 50 μmol m^−2^ s^−1^ (total intensity including green measuring light: 100 μmol m^−2^ s^−1^) for 30 minutes to trigger stomatal closure.

#### *Chrysanthemum* (shade-sun-shade, steady-state, 360 min)

Leaves were acclimated to 50 μmol m^−2^ s^−1^ blue-red light and 50 μmol m^−2^ s^−1^ green imaging light for 60 minutes before image acquisition was started. Image stacks were acquired every minute for 300 minutes. After 60 (120 total) minutes, the light intensity was increased to 1000 μmol m^−2^ s^−1^ by increase of the blue-red light intensity to 950 μmol m^−2^ s^−1^. After another 180 (300 total) minutes, the blue-red light intensity was switched back to 50 μmol m^−2^ s^−1^ (total intensity including green measuring light: 100 μmol m^−2^ s^−1^) for 60 minutes to trigger stomatal closure.

#### *Maize* (growth light - saturating light - growth light, 180 min)

Leaves were acclimated to 425 μmol m^−2^ s^−1^ blue-red light and 75 μmol m^−2^ s^−1^ green imaging light for 60 minutes before image acquisition was started. Image stacks were acquired every minute for 120 minutes. After 30 (90 total) minutes (Period I), the light intensity was increased to 1500 μmol m^−2^ s^−1^ by increase of the blue-red light intensity to 1425 μmol m^−2^ s^−1^ (Period II). After 60 (150 total) minutes the blue-red light intensity was switched back to 425 μmol m^−2^ s^−1^ (total intensity including green measuring light: 500 μmol m^−2^ s^−1^) for 30 minutes to trigger stomatal closure (Period III).

**Table 1.** Glossary.

|  |  |
| --- | --- |
| SS | Steady-state |
| QSS | Quasi-steady-state |
| $A_{0-i}$ | The stomatal aperture at initial SS ($\mu\text{m}^2$) |
| $A_{\text{max}}$ | The maximum stomatal aperture at the final SS after opening ($\mu\text{m}^2$) |
| $\lambda_i$ | The time lag of the opening response (min) |
| $k_i$ | The opening time constant (min), a measure of the rapidity of the opening response to SS |
| Max dAdt-i | Maximum opening speed ($\mu\text{m}^2\text{min}^{-1}$) |
| $A_{0-d}$ | The stomatal aperture at the start of the closing response ($\mu\text{m}^2$) |
| $A_{\text{min-d}}$ | The final SS or QSS after closing ($\mu\text{m}^2$) |
| $\lambda_d$ | The time lag of the closing response (min) |
| $k_d$ | The closing time constant (min), a measure of the rapidity of the closing response to SS |
| Max dAdt-d | Maximum closing speed ($\mu\text{m}^2\text{min}^{-1}$) |
| GCL | Guard Cell Length ($\mu\text{m}$) |
| Voronoi area | The leaf area with the shortest path length to a given stoma ($\mu\text{m}^2$) |
| FWHM | Full width at half maximum |
| LED | Light emitting diode |
| PAR | Photosynthetically active radiation |
| FOV | Field of View |
| WUE | Water-Use-Efficiency |
| NA | Numerical Aperture |
| VPD | Vapor-pressure-deficit |
| CLAHE | Contrast limited adaptive histogram equalization |

**SM Fig 1.**
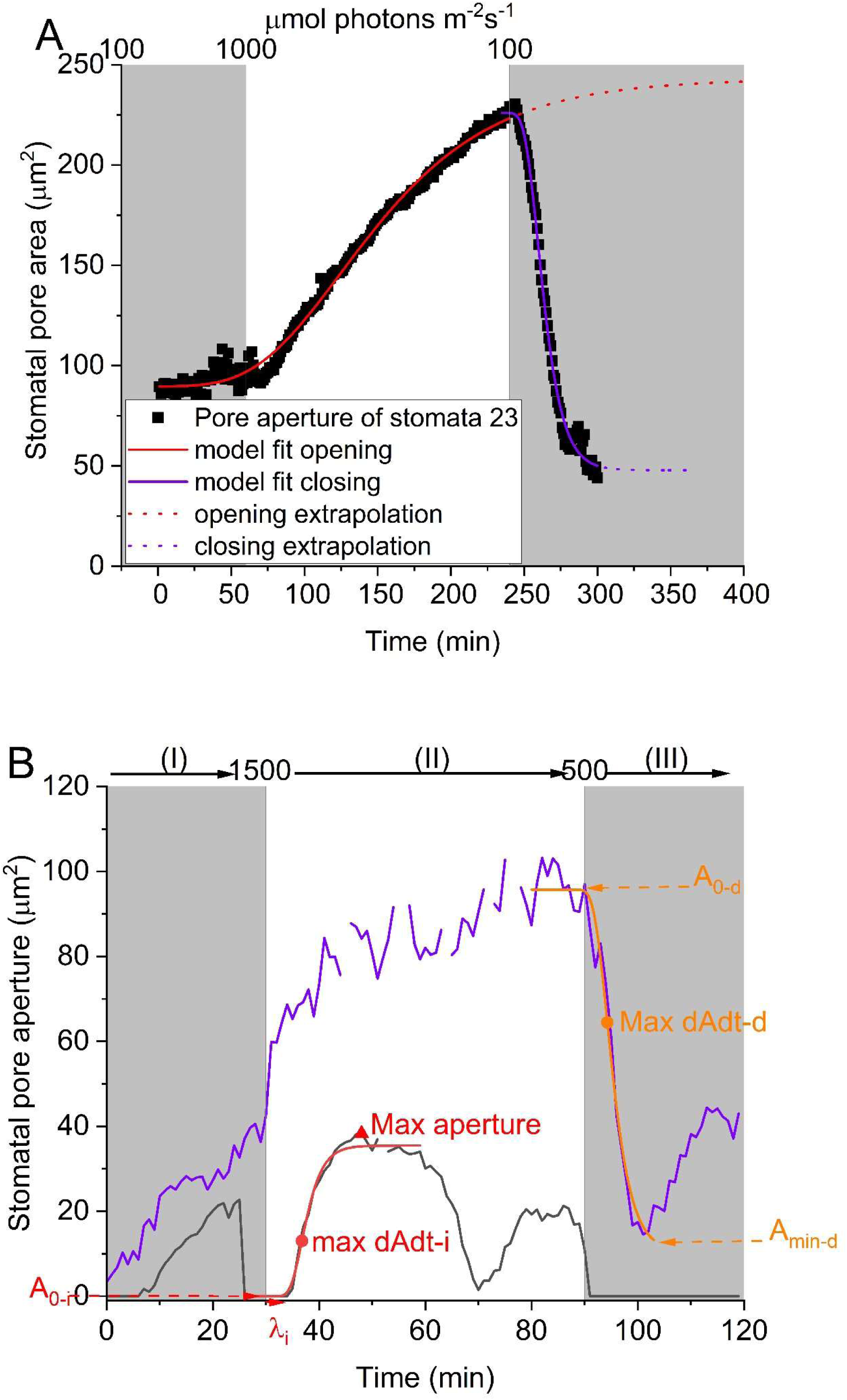
A. Example of stomatal aperture dynamics of a single stoma of Chrysanthemum and the model fit for opening (t=60-240) and closing (t=240-300). Black symbols show raw data, and model fits are shown in red/purple. **B** Examples of maize Stomatal dynamics after a change from growth light intensity (500 µmol photons m^−2^s^−1^) to saturating light intensity (1500 µmol photons m^−2^s^−1^) and back to growth light intensity. Stoma classified as oscillating between open and closed state (I), opening and oscillating (II) and closing to steady-state (III) is shown in black. Stoma classified as open (I), opening towards new steady-state (II) and closing and oscillating (III) is shown in purple. Examples of opening and closing fit, fit parameters, maximum aperture and time position of the maximum opening and closing speeds are indicated (red: opening, orange: closing).

### SM Fig 2 Influence of green imaging light on photosynthesis and stomatal conductance

Green light can reduce the stomatal aperture component driven by blue light at low light intensities (Talbott *et al*., 2002, 2006). To investigate the effect of green imaging light in combination with red and blue light compared to only red-blue light (RB) typically used for photosynthetic induction experiments on stomatal conductance (gsw) and net photosynthesis rate (A), we used our microscopy light sources in an open top gas exchange system (Li-6800, Li-Cor Biosciences). Cuvette size was 3×3 cm^2^, CO_2_ was set to 400 µmol mol⁻¹, flow was 400 µmol s⁻¹, air temperature was 25°C, leaf-to-air VPD was 1.06 ± 0.1 kPa (70% relative air humidity). Data were logged every 30 s with 10 s averaging per timepoint. Temperature of the measurement room was 23±1°C, room RH was 70±5% and room [CO_2_] was ambient. The light spectrum was switched from RB to red-green-blue light (RGB) with approximately the same total light intensity, after reaching a steady state in ~100 or 95% of steady-state gsw at ~1100 µmol photons m^−2^s^−1^ in a Chrysanthemum leaf. Green light intensity was 50 µmol m^−2^s^−1^, and total light intensity was ~100 or ~1100 µmol m^−2^s^−1^.

In low light, switching from RB in the steady state to RGB (G:RB ratio 1:1) quickly reduced A (to a very small extent), whereas gsw was reduced more slowly and strongly (within 34 min (97% of steady-state in RGB). In contrast, no differences were observed in high light, when switching from RGB (G:RB ratio 0.05:1) to RB back to RGB, corroborating previous research (Talbott *et al*., 2002, 2006). In conclusion, the apertures observed in our experiments in low light are smaller than those in the typical RB light used for photosynthetic induction due to the decreasing effect of the green imaging light on stomatal aperture.

**SM Fig. 2.**
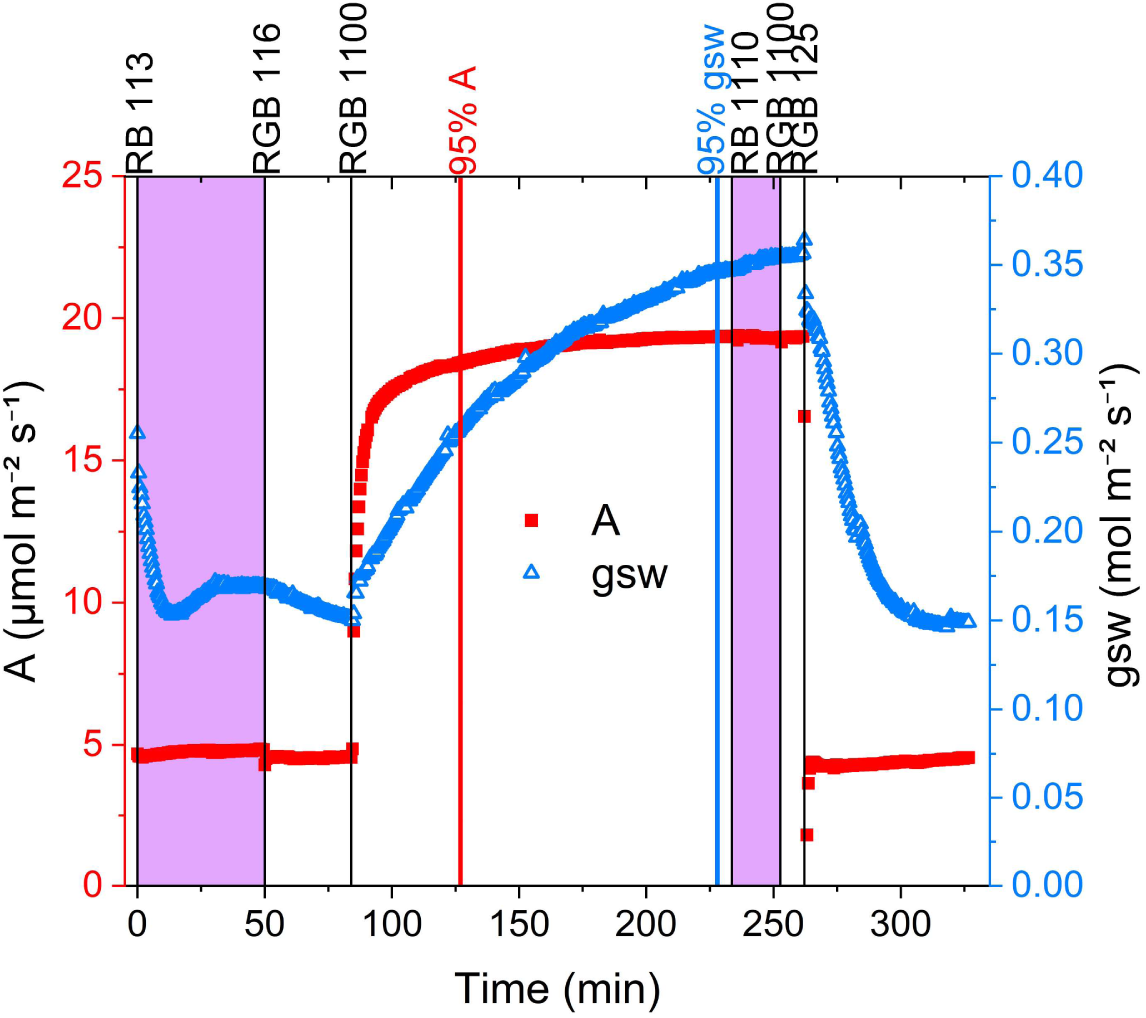
Effect of green imaging light on stomatal conductance and photosynthetic assimilation at 100 and 1100 µmol photons m^2^s^−1^, as measured on a leaf of Chrysanthemum. Stomatal conductance (gsw, blue symbols) and photosynthetic assimilation (A, red symbols) during changes in light spectrum and light intensity are shown. Black vertical lines indicate a change in light spectrum or intensity. Purple areas indicate time period with RB light, white areas indicate time periods with RGB light. Colored vertical lines indicate time-points were 95% of steady-state photosynthetic assimilation (red) or stomatal conductance (blue) are reached.

**SM Fig. 3.**
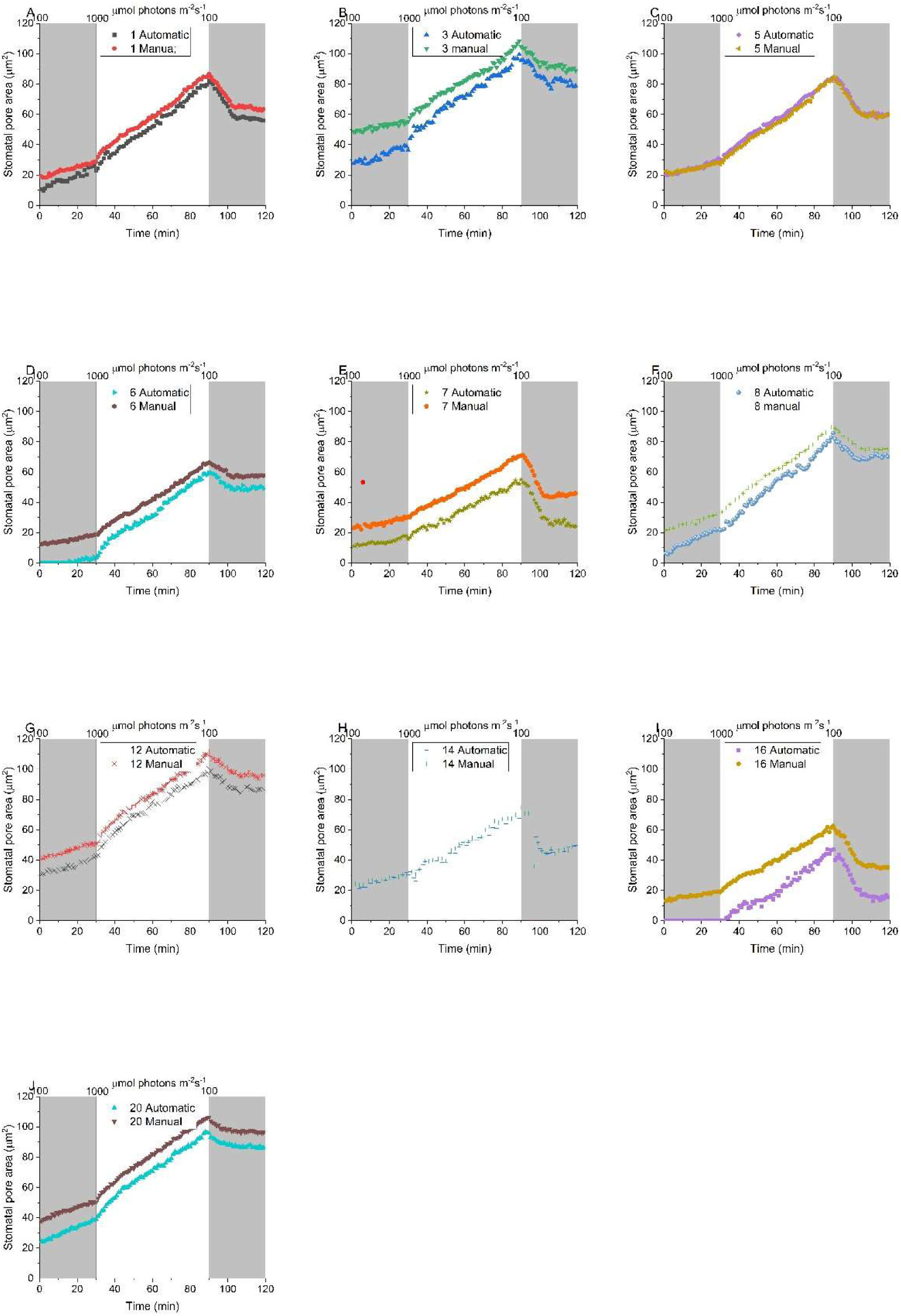
Stomatal aperture dynamics of Chrysanthemum of focus projection images (automatic) and manually determined best single focus images (manual) demonstrating their correlation for 10 individual stomata (A-H)

**SM Fig. 4.**
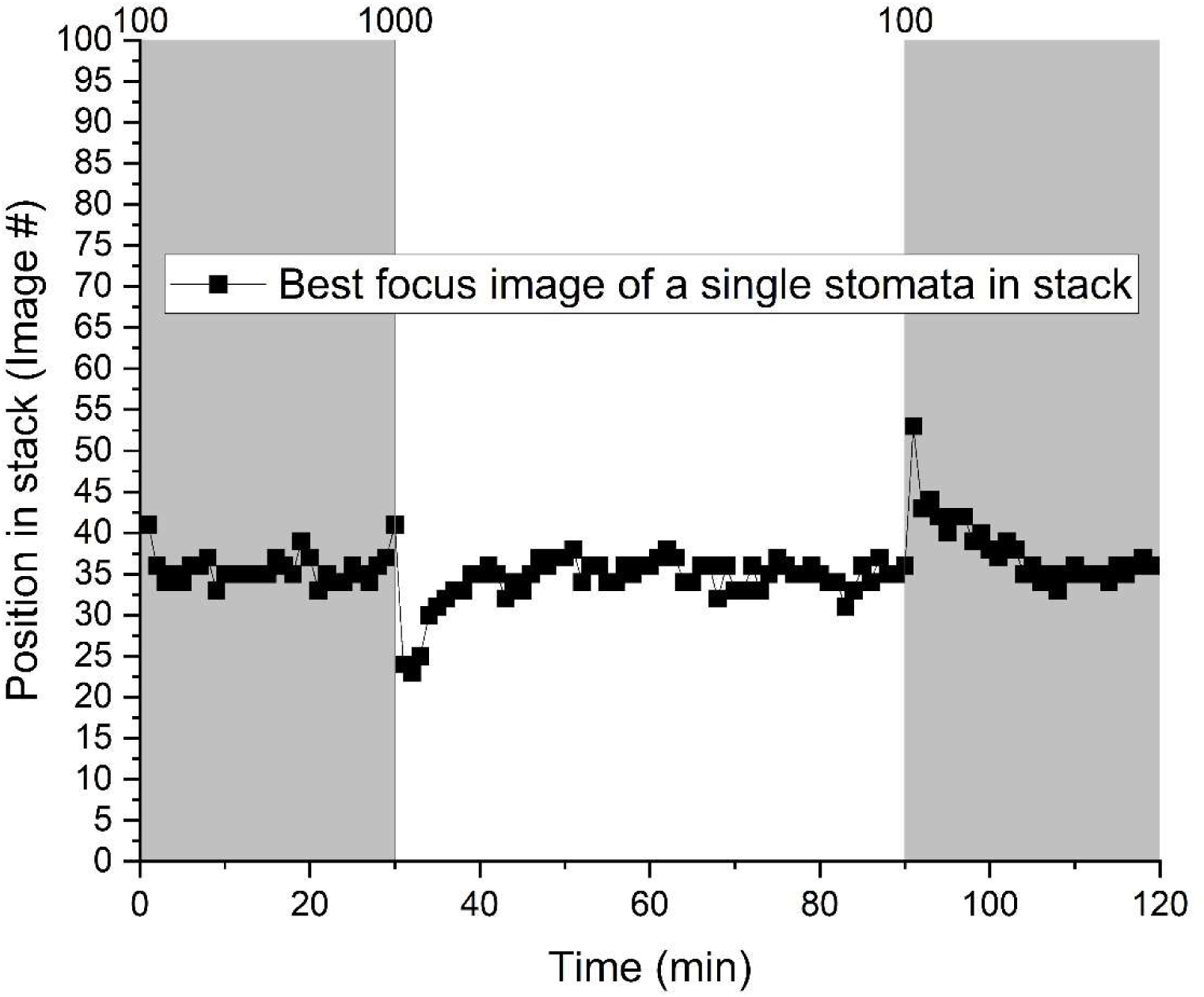
Example of how the position of the best focus for a stoma can change in a stack during the shade-sun-shade experiment in Chrysanthemum due to movement of the leaf surface (Fig. 5).

**SM Fig 5.**
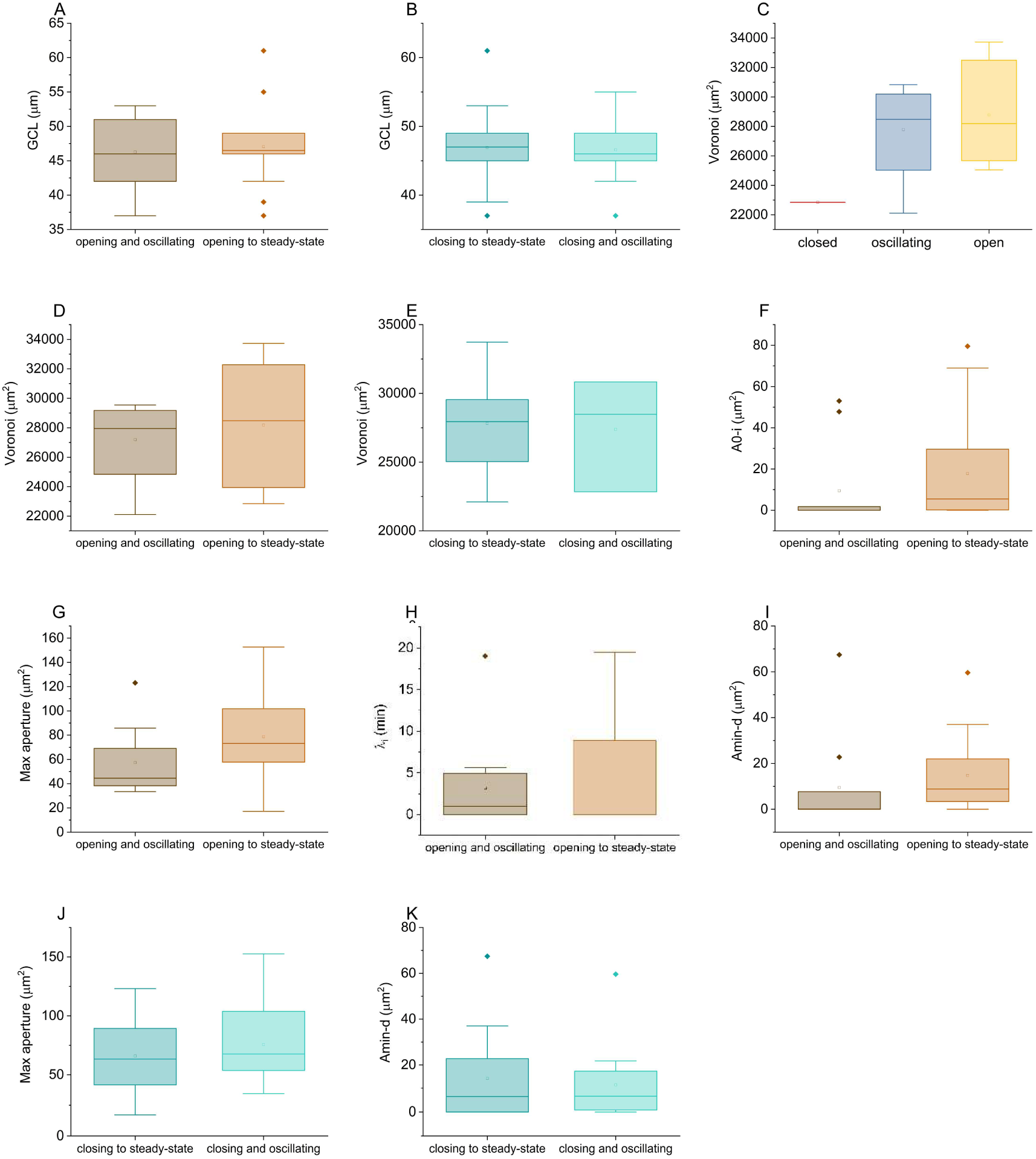
Non-significant differences in comparison of fit and morphological parameters between different categories of Maize stomata tested by ANOVA (AB) or KWANOVA (SW test rejected normality, C-K). Guard Cell length for **A** period II **B** period III. Voronoi area for **C** period I **D** period II **E** period III. **F** Initial, **G** maximum, **I** final aperture and **H** opening lag (λ_i_). **J** maximum and **K** final aperture for period III. Squares represent the average and lines the median (Q2). Box outlines represent the lower (Q1) and upper quartile (Q3) and the whiskers indicate Q0 and Q4.

**SM Fig 6.**
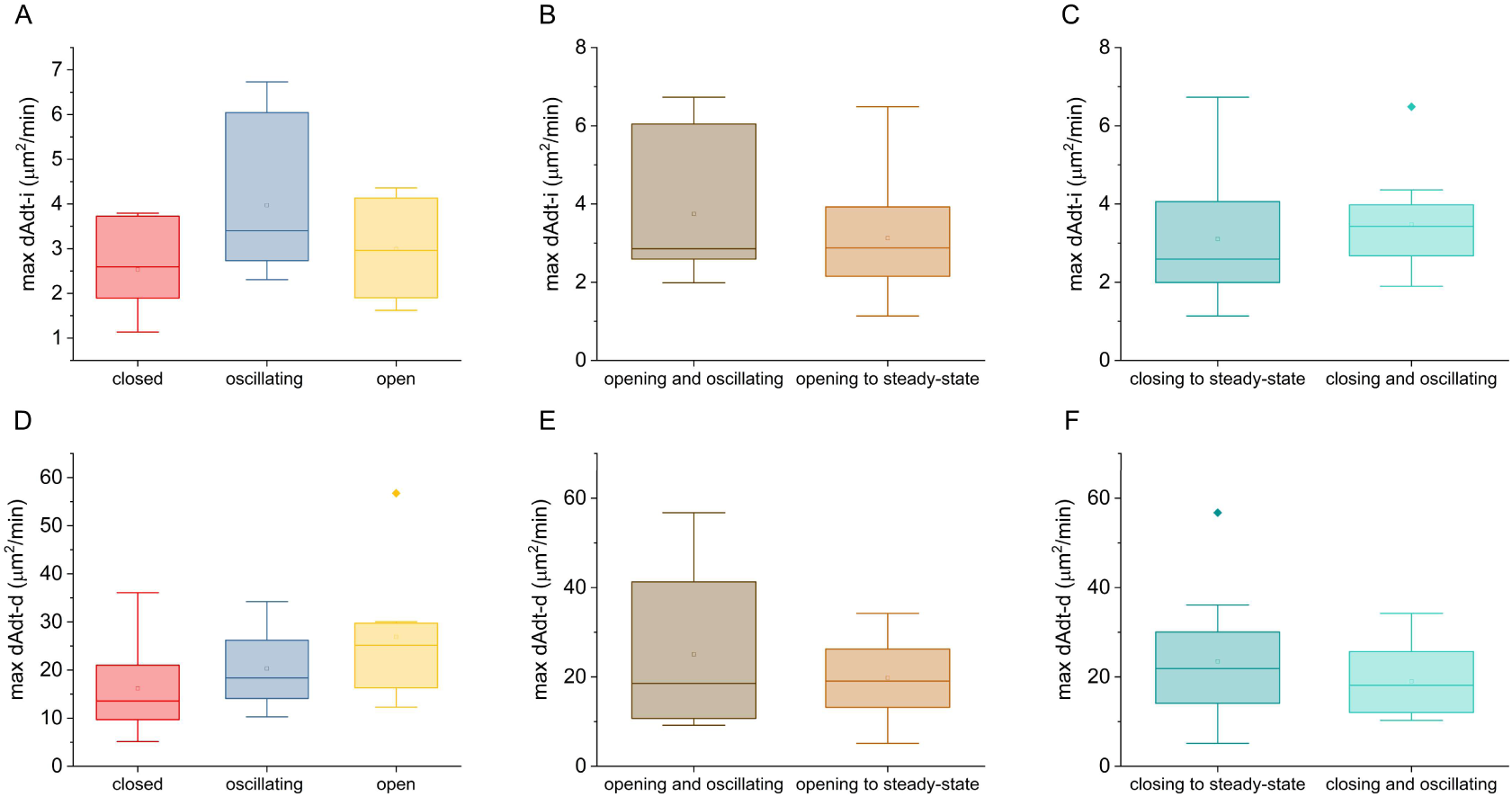
Non-significant differences in comparison of Maize stomatal pore opening (period II) and closing (period III) speeds between different categories tested by KWANOVA (SW test rejected normality). Maximum pore opening speed in period II for category in **A** period I **B** period II **C** period III. Maximum pore closing speed in period III for category in **D** period I **E** period II **F** period III. Squares represent the average and lines the median (Q2). Box outlines represent the lower (Q1) and upper quartile (Q3) and the whiskers indicate Q0 and Q4.

**SM Fig. 7.**
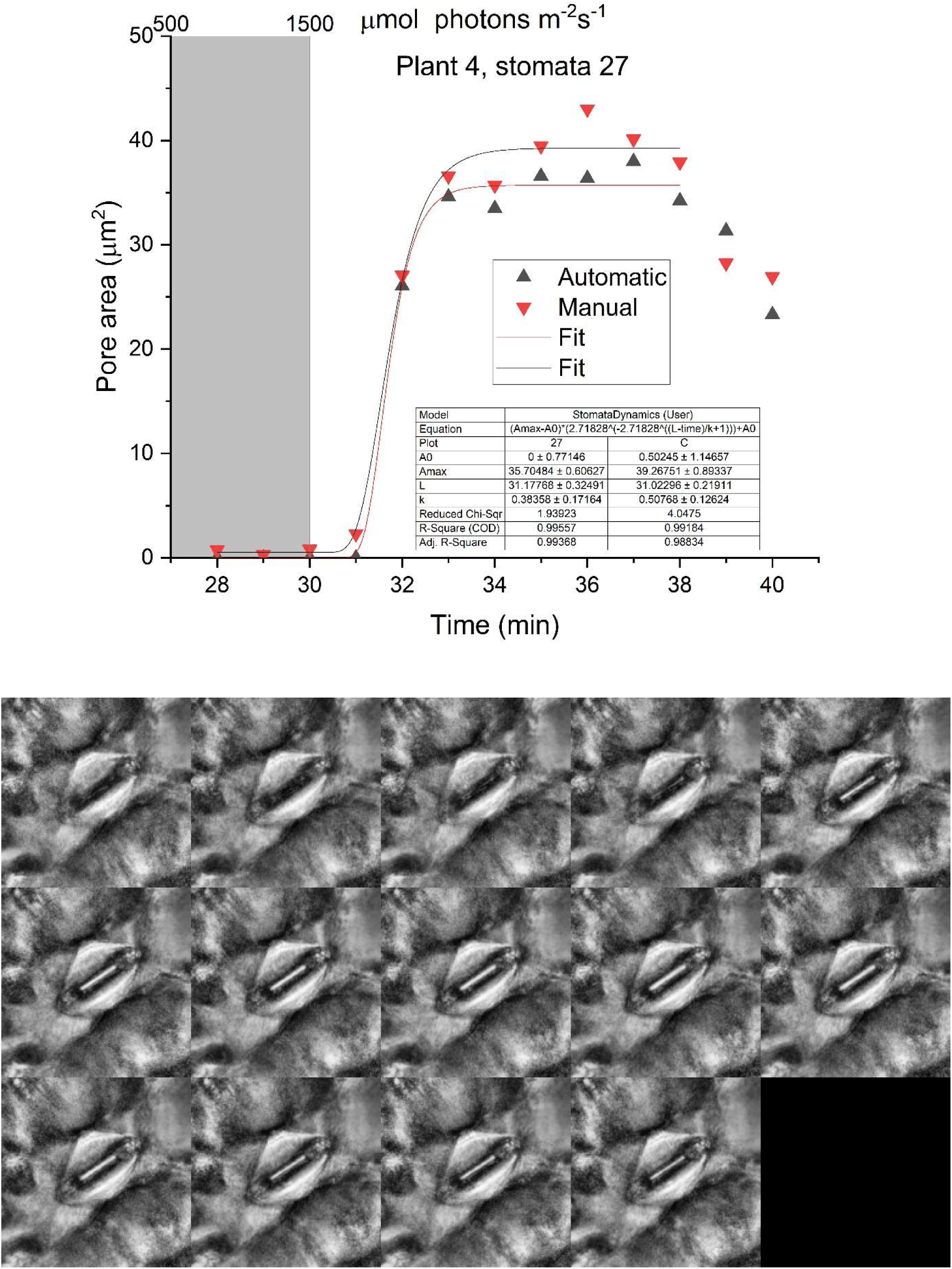
Illustration of large stomatal pore area increase in Maize within a single time-step (<1 min). Timepoints 28 to 41 min are depicted from the best focused single images (Manual, from left to right and top to bottom). Model fits indicate similar results for automatic and manual fits, but both have a large relative error of the fit for the ki parameter.

**SM Fig. 8.**
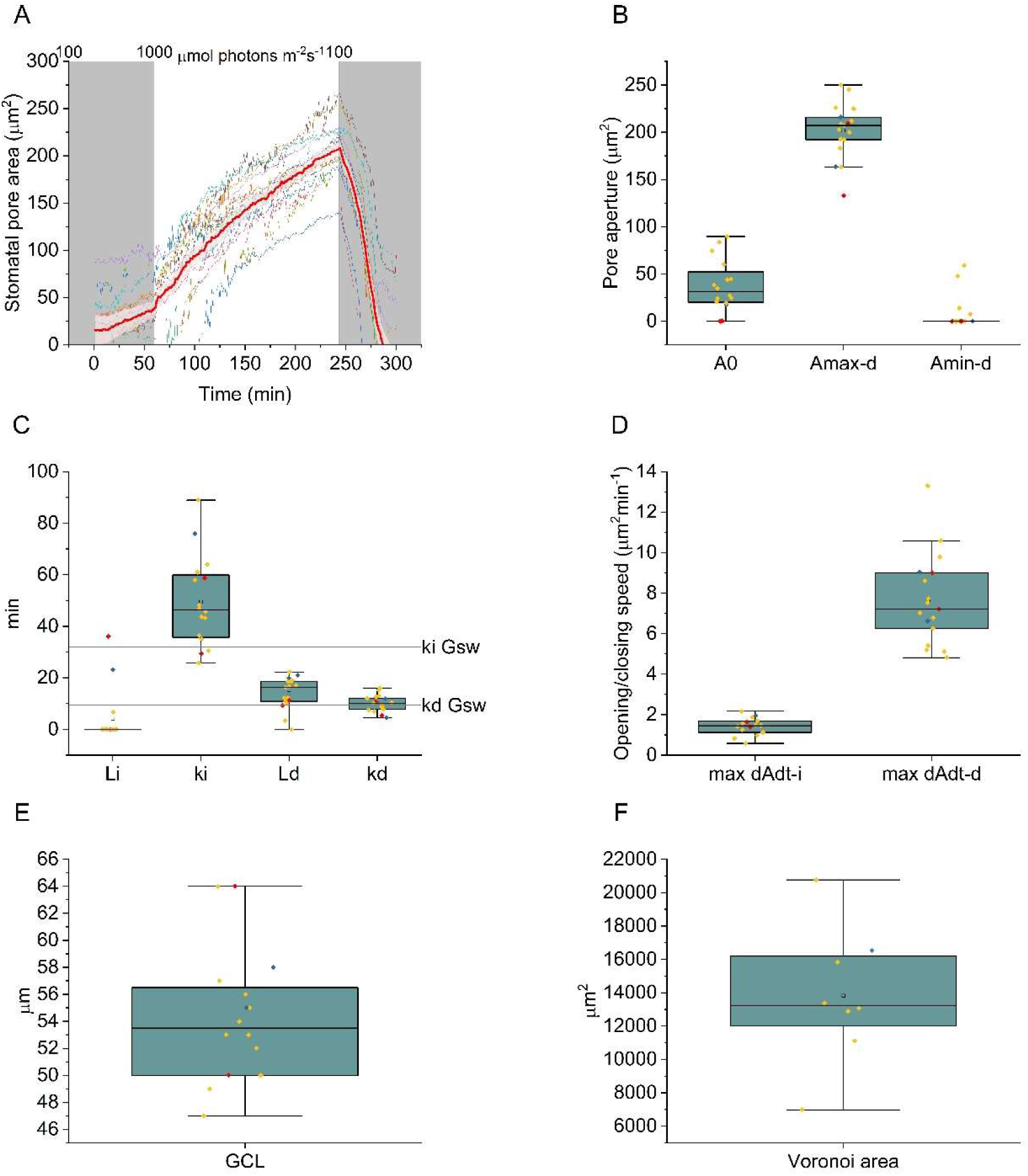
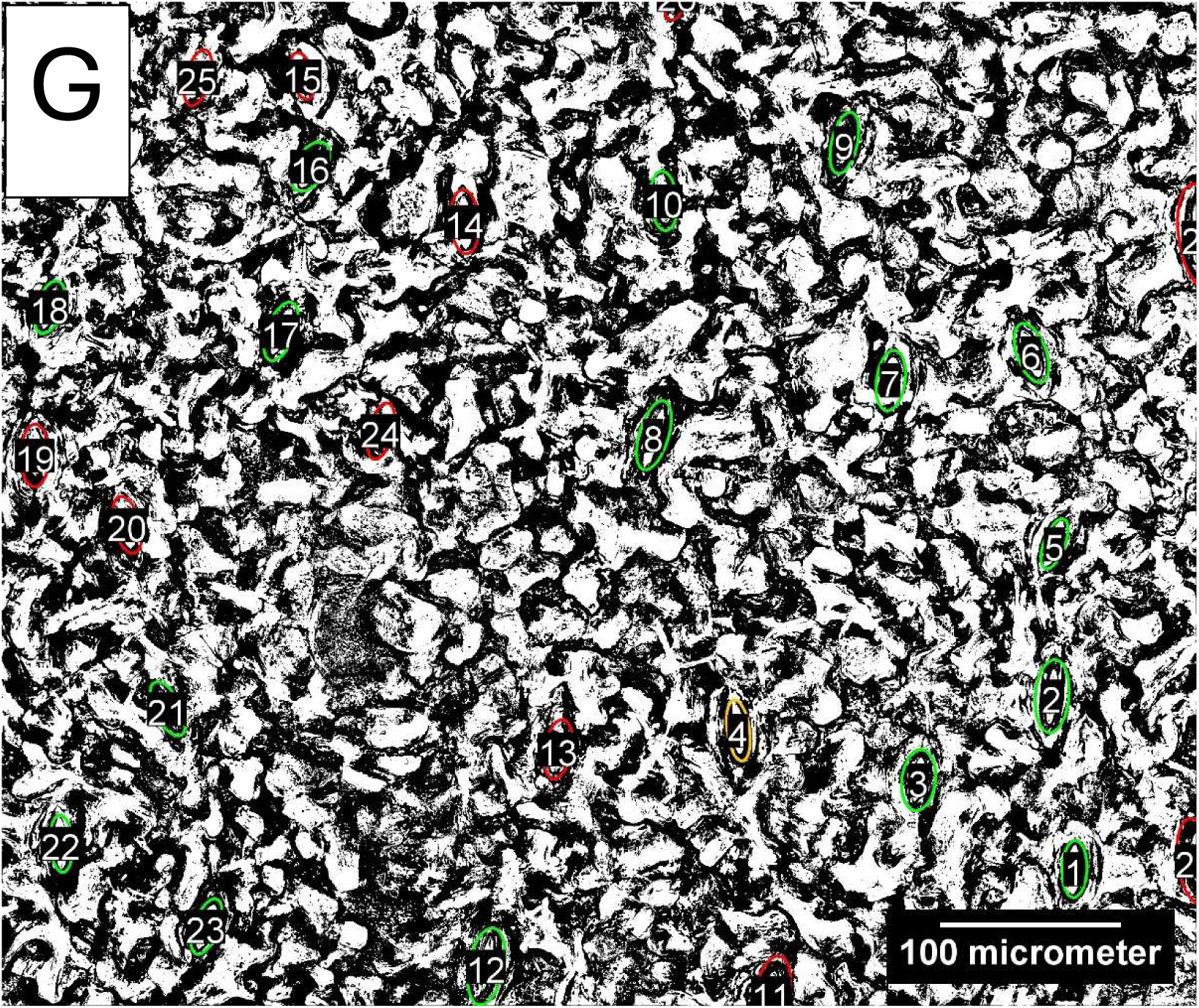
Induction towards steady-state aperture for individual Chrysanthemum stomata **A** traces for all kinetically resolved stomata (17/28) with their average plotted in red. The shaded red area represents the standard error of the mean. Out of the nine unresolved stomata four were on the image edges and three were shaded by a trichome. For the other two there was no clear indication for the poor segmentation of their pores. Out of the 17 resolved stomata two were closed, two were closing and 13 were opening, both without oscillations, during the sixty minutes of low light. In high light all stomata were increasing towards a steady-state without oscillations. In subsequent low light all stomata decreased to a steady-state without oscillations. **B** Initial, maximum and final apertures fitted parameters on the traces in A **C** fitted time constants of opening (k_i_), closing (k_d_) as well as fitted opening (λ_i_) and closing (λ_d_) lag. Horizontal lines represent k_i_ and k_d_ determined in gas exchange (SM Fig. 2). **D** maximum pore opening and closing speed. **E** Guard cell length. **F** Voronoi area **B-F** Color of the data points indicates initial opening (yellow), closing (blue) or closed (red) state of the stoma in low light. **G**. Image of the stomata on the epidermis with the oval shapes enclosing the pore are of the stomata. Green stomata were kinetically resolved, orange only resolved for closing and red unresolved. Squares represent the average and lines the median (Q2). Box outlines represent the lower (Q1) and upper quartile (Q3) and the whiskers indicate Q0 and Q4.

**SM Fig. 9.**
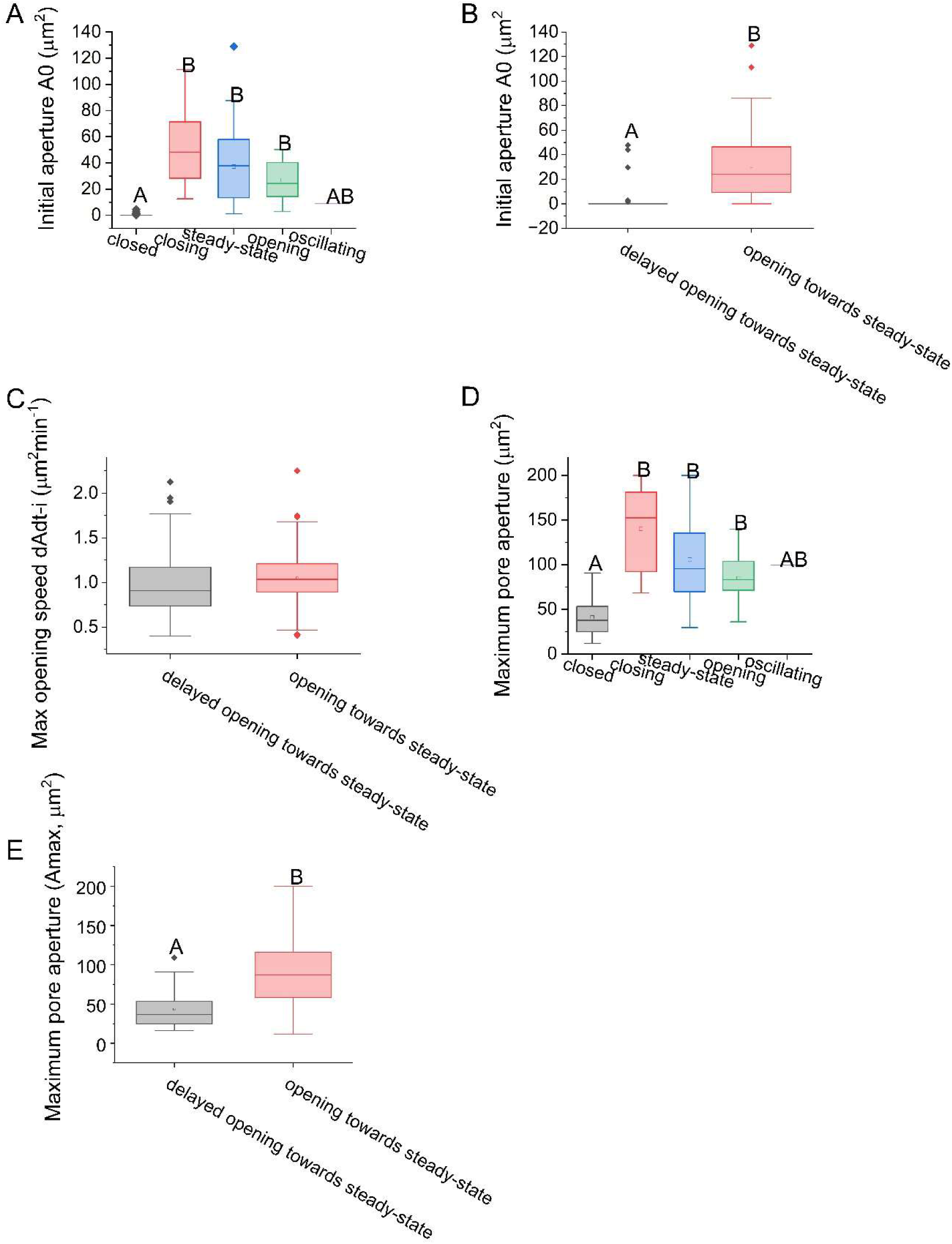
Comparison of kinetic parameters of categorized Chrysanthemum stomata during a shade-sun-shade transition. **A** Initial and **D** maximum pore area of the sun shade phase compared between the kinetic categories of the initial shade phase. Number of stomata per category: 47 closed, 25 open in a steady-state, 27 opening, 21 closing and one stoma was open but oscillating. **B** Initial pore aperture, **C** maximum pore opening speed and **E** Maximum pore aperture in the sun phase compared between the kinetic categories of the sun phase. Number of stomata per category: 87 opening towards steady-state. 37 Delayed opening towards steady-state. Squares represent the average and lines the median (Q2. Box outlines represent the lower (Q1) and upper quartile (Q3) and the whiskers indicate Q0 and Q4.

SM Movie 1 Example of a processed z-stack (100images) of a single timepoint in the shade-sun-shade experiment of Chrysanthemum. Stomatal positions are indicated by red ovals.

SM movie 2 Example of the processed time stack (video) of a complete experiment (119 focus projections of the z-stacks of each timepoint) in the shade-sun-shade experiment of Chrysanthemum. Stomatal positions are indicated by red ovals.

SM video 3 Example of the processed time stack (video) of a complete shade-sun-shade experiment of Chrysanthemum, zoomed in on a single stoma.

SM video 4 The same video of 3 after thresholding.

SM video 5 The same video as 3 and 4 after segmentation.

